# Notch Signaling Drives Pro-Regenerative and Migratory Traits in Glandular Stem/Progenitor cells

**DOI:** 10.1101/2024.11.27.625738

**Authors:** Davide Cinat, Rufina Maturi, Jeremy P. Gunawan, Anne L Jellema-de Bruin, Laura Kracht, Paola Serrano Martinez, Yi Wu, Abel Soto-Gamez, Marc-Jan van Goethem, Inge R. Holtman, Sarah Pringle, Lara Barazzuol, Rob P. Coppes

## Abstract

Organoid models have advanced our understanding of adult stem/progenitor cell dynamics and function. However, uncovering the regulatory mechanisms of scarce and often quiescent stem cells in organs like the salivary glands remains challenging. Using single-cell RNA sequencing and bulk ATAC and RNA-sequencing analysis, we conducted in-depth profiling of the cellular populations and key signaling pathways characterizing a mouse submandibular salivary gland organoid (mSGO) model at different temporal stages and in response to radiation damage. We identified *Sox9-* and *Itgb1-*expressing cells as the most primitive adult stem/progenitor populations and uncover novel stemness and migratory roles for *Cd44*-expressing cells. Moreover, we revealed that Notch signaling is essential for maintaining self-renewal and migration potential of these stem/progenitor cells post-irradiation. Extending these findings to patient-derived mSGOs, as well as murine and patient-derived mammary and thyroid gland organoids, we confirmed Notch signaling as a conserved regulator of stem/progenitor cell function under migrative and regenerative conditions.

## Introduction

Adult stem/progenitor cells in glandular tissues, like the salivary glands, are scarce and often residing in a quiescent state, making their identification challenging^1^. Moreover, their regenerative capacities in response to damage have been shown to be heavily influenced by the microenvironment, impacting self-renewal and differentiation abilities^2,3^. Given their critical role in maintaining tissue homeostasis^4^ and the demand for personalized and regenerative therapies^5,6^, there has been growing interest in characterizing their identity and regulatory pathways^4,5^.

Salivary glands are exocrine organs composed of serous and mucous saliva-secreting acinar cells, myoepithelial cells, and a network of intercalated, luminal and basal duct cells^1^. Among these, the basal layer of the ductal compartment is believed to host putative stem/progenitor cells^1,7^. Although organoids have proven to be a valuable model for characterizing salivary gland stem/progenitor cells^8,9^, a comprehensive understanding of the cells constituting this organoid model and the signaling pathways regulating their regenerative functions is still lacking.

Radiotherapy is one of the most common therapeutic modalities for treating head and neck cancer^10^. However, due to the close proximity to the radiation site, salivary glands often suffer collateral damage after treatment^10,11^, resulting in the loss of stem/progenitor cell populations and subsequent impairment of tissue regeneration^11,12^. This can result in side effects, such as radiation-induced hyposalivation and related xerostomia, significantly impairing patients’ quality of life^13^. Salivary gland-derived organoids have played a salient role in elucidating the molecular mechanisms underlying the response to radiation-induced DNA damage^9,14,15^; however, the impact of irradiation on specific stem/progenitor populations important for regeneration is still unknown.

The Notch signaling pathway is one of the key regulators of stem cell fate decision and self-renewal capacity in several tissues^16^. The canonical Notch signaling cascade consists of a highly conserved cell-cell communication mechanism, triggered by the interaction between the ligand expressed on one cell, and the corresponding Notch receptor on a neighboring cell^16,17^. Subsequent to ligand-receptor binding, cleavage of the intracellular domain of the Notch (NICD) receptor by γ-secretase allows NCID translocation into the nucleus and transcription of its target genes^16,17^. Although the Notch pathway has been extensively studied in various tissues and organs^18–21^, its role in regulating glandular stem/progenitor cells remains to be fully elucidated.

In this study, we employed single-cell RNA sequencing (scRNA-seq), bulk RNA and ATAC-sequencing to explore the dynamic development of a mSGO model across two different temporal stages, as well as following photon and proton irradiation, two clinically relevant radiation types for the treatment of head and neck cancer. We identified novel adult stem/progenitor features of *Sox9-*, *Itgb1-,* and *Cd44-*expressing cells, highlighting their primitive characteristics relative to the other populations. Furthermore, we revealed for the first time that Notch signaling plays a crucial role in regulating the regenerative capacity of both mouse and human submandibular salivary gland organoid-derived stem/progenitor cells after radiation damage, as well as in human mammary and thyroid gland organoids.

## Results

### scRNA-seq of mSGOs show distinct populations resembling salivary gland tissue composition

To obtain a deeper understanding of the populations constituting salivary gland organoids, adult mouse submandibular salivary gland tissue was dissociated, and single cells were placed and cultured in Matrigel^22^. After one passage, mSGOs were collected on day 7 and 11 of culturing and processed for scRNA-seq and subsequence analysis (Figure 1A). These time points were chosen as they represent the optimal stages for obtaining fully formed salivary gland organoids^23,24^ for studying the regenerative response of salivary gland stem/progenitor cells *in vitro*^15,25^. Upon filtering and exclusion of low-quality cells (see methods), 7.510 cells from 7-day (Figure 1B) and 11.248 cells from 11-day organoids (Figure 1C) were mapped. Unsupervised clustering was performed, resulting in 6 distinct clusters for 7-day organoids (Figure 1D) and 8 distinct clusters for 11-day organoids (Figure 1E). Based on established markers^24,26,27^, we identified two populations of potential stem/progenitor cells, namely: stem/progenitor cell I (*Cd164, Sox9, Cd24a*) and stem/progenitor cells II (*Itgb1, Cd44, Trp63*); basal duct cells (*Krt14*, *Krt5*); luminal duct cells (*Krt8*, *Krt7/17, Cldn4, Clic1*); and, cycling cells (*Mki67, Top2a*) in both 7-day and 11-day mSGOs (Figures 1D-G). 7-day mSGOs were mostly composed of basal duct and stem/progenitor cells (Figure S1A), which constitute the most plastic cells in the salivary gland^1^, indicative of a premature phenotype. In contrast, analysis of 11-day mSGOs revealed a shift towards a more differentiated phenotype including an increased number of luminal duct cells (Figure S1B), pro-acinar cells (*Chrm3, Alcam, Clcn3*)^28–30^ and myoepithelial-like cells (*Trp63, Lgals7, Csrp1, Sparc*)^31–33^ (Figures 1E, 1G), aligning with the finding of our previous published study and closely resembling the *in vivo* tissue composition^24^. Salivary gland stem/progenitor cells are believed to primarily reside within the ductal compartment of adult salivary glands^11^. To test this hypothesis and compare the expression profiles of mSGOs with those of adult tissue, we performed scRNA-seq analysis of freshly isolated murine submandibular salivary glands and compared it with the 7-day mSGO dataset. After filtering out low-quality cells, 2.519 cells from the salivary gland tissue samples were mapped (Figure S1C, S1D). Interestingly, a subpopulation of duct cells extrapolated from the scRNA-seq tissue dataset (Figure 1H and S1C, S1D) showed a strong expression of genes associated with both mSGO-stem/progenitor cells, such as *Cd164, Sox9, Cd24a*, *Itgb1* and *Cd44* (Figure 1I and S1E). This suggests the potential presence of a stem cell-like population within the ducts of the salivary glands. Overall, these data confirm an enrichment of salivary gland stem/progenitor cells within younger mSGOs and the presence of more differentiated cell types in older mSGOs, resembling the originating salivary gland tissue composition.

**Figure 1.**
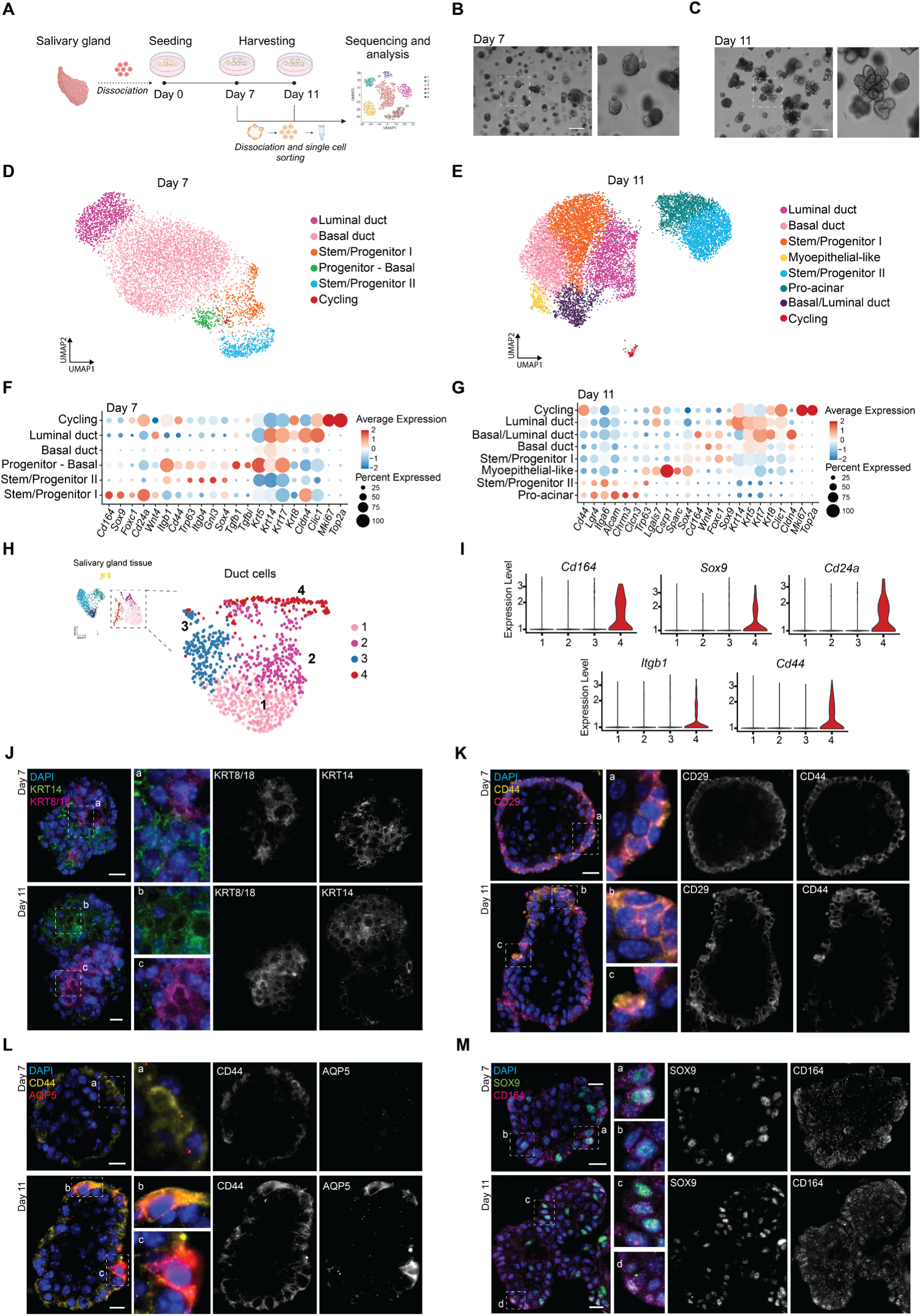
scRNA-seq of mSGOs reveals distinct cell populations and an enrichment of stem/progenitor cells. (**A**) Schematic representation of the scRNA-seq experiment. (**B**) Representative image of 7-day mSGOs. Scale bar, 100 µm. (**C**) Representative image of 11-day mSGOs. Scale bar, 100 µm. (**D**) UMAP of the 7-day mSGO dataset showing the main cell populations. (**E**) UMAP of the 11-day mSGO dataset showing the main cell populations. (**F**) Dot plot showing cell type marker genes of the 7-day-old mSGO dataset. (**G**) Dot plot showing representative marker genes of the 11-day mSGO dataset. (**H**) UMAP of a subpopulation of duct cells extrapolated from the scRNA-seq dataset of salivary gland tissue showing the main cell sub-clusters. The entire UMAP of salivary gland tissue is shown in Figure S1C. (**I**) Violin plots showing the expression of *Cd164*, *Sox9, Cd24a, Itgb1* and *Cd44* across the different Duct sub-clusters. Colors are the same as in Figure 1H. (**J**) Representative images of immunofluorescence staining of 7-day and 11-day mSGOs showing the expression of KRT14 and KRT8/18. Scale bar, 10 µm. (**K**) Representative images of immunofluorescence staining of 7-day and 11-day mSGOs showing the expression of CD29 and CD44. Scale bar, 10 µm. (**L**) Representative images of immunofluorescence staining of 7-day and 11-day mSGOs showing the expression of CD44 and AQP5. Scale bar, 10 µm. (**M**) Representative images of immunofluorescence staining of 7-day and 11-day mSGOs showing the expression of SOX9 and CD164. Scale bar, 10 µm.

In line with the scRNA-seq data, immunofluorescence staining of organoid sections showed the presence of two distinct populations of duct cells: luminal duct cells identified by KRT8/18 and basal duct cells identified by KRT14 (Figure 1J). Moreover, we identified cells positive for CD29 (*Itgb1*) and CD44 constituting a population of stem/progenitor cells (Figure 1K). Interestingly, AQP5, a well-known pro-acinar and acinar marker, was found to be expressed by some CD44 positive cells, especially in 11-day mSGOs (Figure 1L). This aligned with our scRNA-seq data (Figure 1E), which suggested a potential role for the *Itgb1/Cd44-* expressing stem/progenitor population as acinar precursor cells. *Cd164*, a gene encoding a sielomucin protein described as a novel human stem cell marker^34^, was highly expressed by the *Sox9+* stem/progenitor cluster (Figure 1F, 1G). Immunofluorescence staining confirmed this finding, identifying a co-expression of SOX9 and CD164 at the protein level (Figure 1M).

Together, our data establish the existence of distinct cell populations within our mSGOs and provide new insights into salivary gland stem/progenitor cell populations and their markers.

### Salivary gland stem/progenitor cells are functional in mSGOs

Based on known stem/progenitor cell markers, scRNA-seq analysis revealed the presence of two salivary gland stem/progenitor populations within mSGOs (Figures 1D, 1E). Notably, genes identifying these populations were strongly associated with stem cell-related gene ontology terms, such as embryonic and tissue development-related biological processes (Figure S1F), indicating stem/progenitor cell features. To confirm these findings, we performed ATAC-sequencing (ATAC-seq) of 7-day mSGOs, which exhibited a more premature phenotype compared to 11-day mSGOs. This analysis revealed an enrichment of several motifs associated with transcription factors strongly linked to embryonic development, tissue development, differentiation and morphogenesis-related processes (Figure S2A, S2B). Specifically, motifs associated to KLF, TEAD, RUNX, SOX and FOX, which are known regulators of embryonic and stem cell-related processes^35–39^, were highly enriched in the stem/progenitor clusters (Figure 2A), suggesting heightened stem cell features compared to the other populations. To further support this notion, we integrated the 7-day and 11-day mSGO datasets (Figure 2B) and performed unbiased trajectory and pseudotime analyses (Figures 2C and S2D). Notably, the pseudotime trajectory originated from a cluster of cells of 7-day mSGOs expressing higher levels of stem/progenitor markers including *Sox9, Itgb1, Cd24a and Trp63* (Figures 2D and S2E). The expression of these genes decreased along the pseudotime, in contrast to more differentiated markers such as *Krt7*, *Alcam, Chrm3 and Clcn3*, which increased towards the end of the trajectory in 11-day mSGOs (Figures 2D and S2E). In line with this, bulk-RNA sequencing of 5-, 7– and 11-day mSGOs showed increased expression of differentiated markers over time (Figure 2E), indicating maturation and a more differentiated phenotype at day 11. Together, these data indicate an enrichment of stem/progenitor cells in 7-day mSGOs and highlight *Sox9-* and *Itgb1*-expressing cells as the most primitive cell types within our mSGOs.

**Figure 2.**
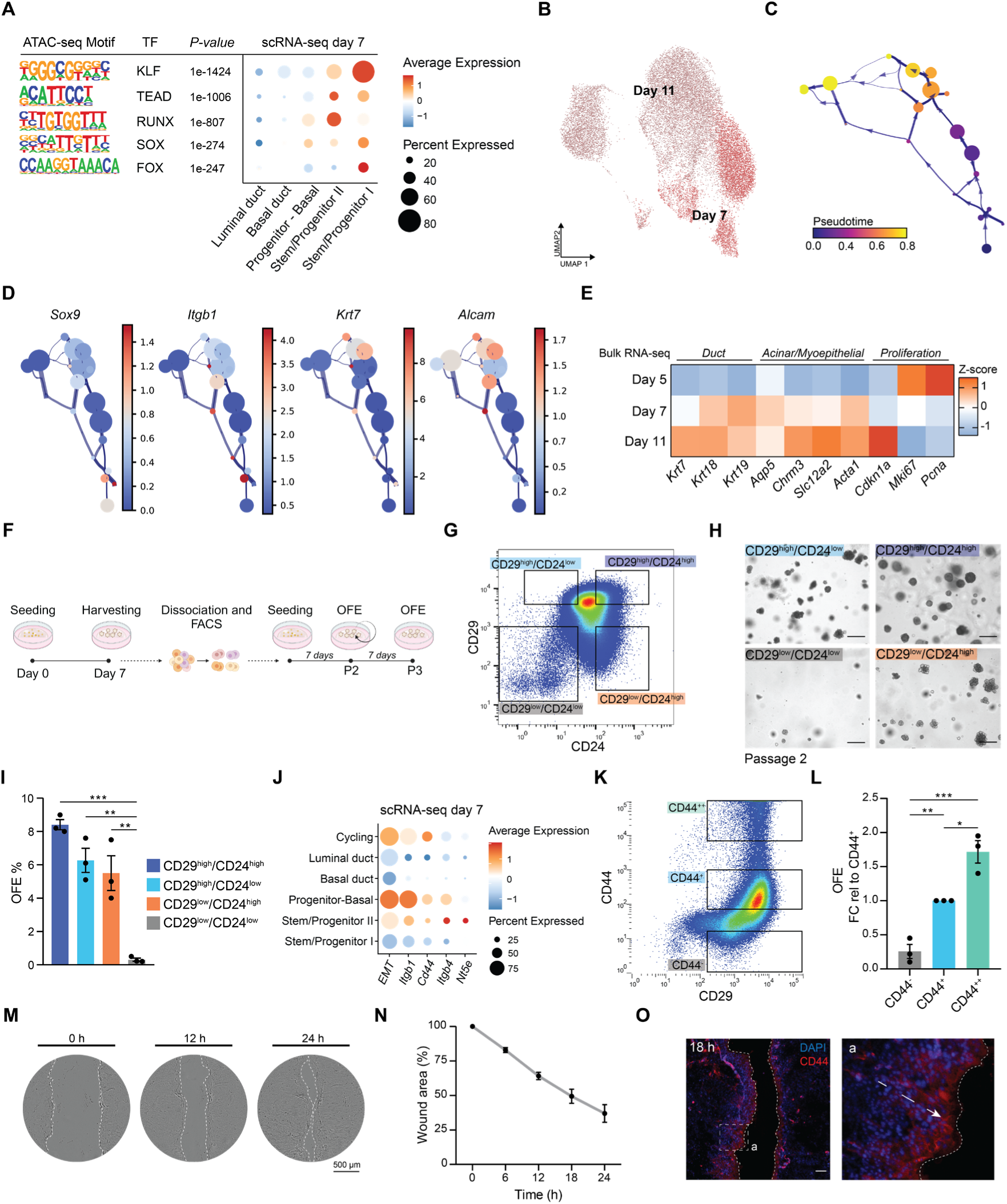
*Sox9-, Itgb1-* and *CD44-* expressing cells display stem/progenitor features. (**A**) Table showing the ATAC-seq motifs and relative *P*-values of KLF, TEAD, RUNX, SOX and FOX transcription factor (TF) families. Expression of the genes associated to each motif was then extrapolated from the 7-day mSGOs dataset. (**B**) UMAP of the 7-day and 11-day mSGO-integrated dataset. (**C**) Trajectory plot of the 7-day and 11-day mSGO-integrated dataset showing the pseudotime course. Trajectory and cluster distribution is shown in Figure S2D. (**D**) Trajectory plot of the 7-day and 11-day mSGO-integrated dataset showing the expression of *Sox9, Itgb1, Krt7* and *Alcam*. (**E**) Heatmap showing the expression of Duct, Acinar, Myoepithelial and Proliferation-related genes extrapolated from the bulk-RNA sequencing datasets of 5-day, 7-day and 11-day mSGOs. Data is shown as Z-score. (**F**) Schematic representation of the self-renewal assay following FACS of 7-day mSGOs. (**G**) Representative FACS plot showing the gating strategy. Unstained and back gating is shown in Figure S3A, S3B. (**H**) Representative images of sorted cells after 1 week in culture (P2). Scale bar, 100 µm. (**I**) Organoid quantification of sorted cells after 1 week in culture (P2) shown as organoid formation efficiency (OFE%) (means ± s.e.m; n = 3 animals/condition). One-way ANOVA, post-hoc Tukey’s test. ** *p* < 0.01, *** *p* < 0.001. (**J**) Dot plot showing mesenchymal-like markers and EMT features (GSEA Mouse Gene Set: Hallmark Epithelial Mesenchymal Transition) in the 7-day mSGO dataset. (**K**) Representative FACS plot showing the gating strategy. (**L**) Organoid quantification of sorted cells after 1 week in culture shown as organoid formation efficiency (OFE). Data is shown as FC relative to CD44^+^ (means ± s.e.m; n = 3 animals/condition). One-way ANOVA, post-hoc Tukey’s test. * p < 0.05, ** p < 0.01, *** p < 0.001. (**M**) Representative images of salivary gland cells at 0, 12 and 24 h after wound generation. Dotted line shows wound borders. (**N**) Quantification of the wound area over time. Data is relative to 0 h (100%). (**O**) Representative images of immunofluorescence staining of salivary glands cells 18 h after wound generation showing the expression of CD44. Scale bar, 100 µm.

ScRNA-seq data revealed that *Itgb1* (CD29) was highly upregulated in the stem/progenitor cell cluster II, whereas *Cd24a* (CD24), previously associated with a potential population of salivary gland stem cells^23^, was highly enriched in the stem/progenitor I population (Figures 1F and S3A). To validate the stem cell properties of these putative stem/progenitor cells identified in the mSGO datasets, we used fluorescence-activated cell sorting (FACS). Based on the surface markers highly expressed in our scRNA-seq data, we sorted the stem/progenitor populations from 7-day mSGOs. We next assessed their ability to form organoids and self-renew *in vitro* (Figure 2F and S3B, S3C). When sorted cell populations were cultured in Matrigel, CD29^high^/CD24^low^ and CD29^low^/CD24^high^ expressing-cells (Figure 2G) demonstrated a significantly greater capacity to form organoids (Figures 2H, 2I) and ability to self-renew, as indicated by a higher secondary OFE (Figures S3D, S3E), compared to the CD29^low^/CD24^low^-population, thus confirming their stem cell properties. Additionally, we identified and sorted a CD29^high^/CD24^high^ population (Figure 2G) that also exhibited high self-renewal capacity (Figures 2I and S3E), suggesting the presence of a double-positive stem/progenitor population. These findings corroborate the presence and functional capabilities of salivary gland stem/progenitor cells within our mSGOs, substantiating our previous findings^23^.

Further scRNA-seq analysis revealed an upregulation of genes associated with mesenchymal features, including *Cd44, Itgb4, Nt5e*^40^, and a set of epithelial/mesenchymal (E/M)-related genes (EMT), in the *Itgb1-*expressing populations (Figure 2J). Particularly, *Cd44* has been described as a key feature in E/M states, playing an essential role in tumor progression by promoting both stemness and migration of cancer stem cells^41^, and serving as an important marker for hematopoietic stem cells^42^. Given its high expression in the stem/progenitor II and cycling populations (Figures 1F, 1G), we investigated whether *Cd44^high^*-expressing cells exhibited enhanced stem/progenitor cell properties and increased migratory potential. FACS analysis of CD29^high^ cells showed different levels of CD44 (Figure 2K), with higher CD44 expression (CD44^++^) correlating with increased OFE, suggesting enhanced stemness features (Figure 2L). To assess the potential migratory behavior of these cells, we conducted a wound-healing assay in 2D salivary gland cultures derived from dissociated mSGOs, with serum starvation to limit cell proliferation. Notably, these salivary gland cultures, containing the primary cell populations found in the 3D organoid model (Figure S3F), exhibited a clear migration potential and the ability to fill the wound over time (Figures 2M, 2N). Moreover, CD44-expressing cells accumulated prominently at the wound’s migratory front, migrating from the outer regions towards the wound center (Figure 2O). These findings, along with the co-expression of AQP5 by some CD44-expressing cells in 11-day mSGOs (Figure 1L), suggest that CD44 may identify a previously unrecognized salivary gland cell population with both stemness and migratory properties.

### Irradiation-surviving stem/progenitor cells show increased activity and Notch signaling

To gain a deeper understanding of the effects of radiation on the regeneration capabilities of stem/progenitor populations within our mSGOs, organoids were irradiated with photon and proton, two clinically relevant radiation modalities, followed by scRNA-seq analysis. mSGOs were irradiated at day 5 with 7 Gy photon or proton, a dose previously observed to induce a significant cytotoxic and regenerative response, as well as mechanistic changes consistent with *in vivo* models^9,15^, and collected at day 7 for scRNA-seq and analysis (Figure 3A). Upon exclusion of low-quality cells (see methods), the datasets from control, photon– and proton-irradiated samples were merged (Figure 3B) and unsupervised clustering was performed (Figures 3C, S4A). Cluster annotation was executed by using the previously identified markers (Figure 1F) and the presence of similar populations compared to control samples were identified (Figures S4B, S4C). To detect gene expression differences between control, photon and proton irradiation, we performed differential gene expression (DEG) analysis and gene set enrichment analysis (GSEA) for each cluster individually. In line with our previous works^14,15^, GSEA analysis revealed an overall upregulation of immune and mitochondrial stress-related gene ontology (GO) terms after irradiation with both photons and protons (Figure 3D). However, the stem/progenitor and basal duct cells showed an enrichment in processes related to proliferation, development and tissue morphogenesis. These processes include GO terms such as gland development, tube morphogenesis, tube development and cell proliferation (Figure 3D), suggesting an upregulation of stem cell and regenerative features after irradiation. Notably, *Hmga2* and genes members of the Notch signaling pathway, such as *Notch1* and *Hes1,* emerged as the most enriched genes associated with these stem cell-related biological processes (Figure 3E), suggesting their potential role in the maintenance of the surviving stem/progenitor cells after irradiation.

**Figure 3.**
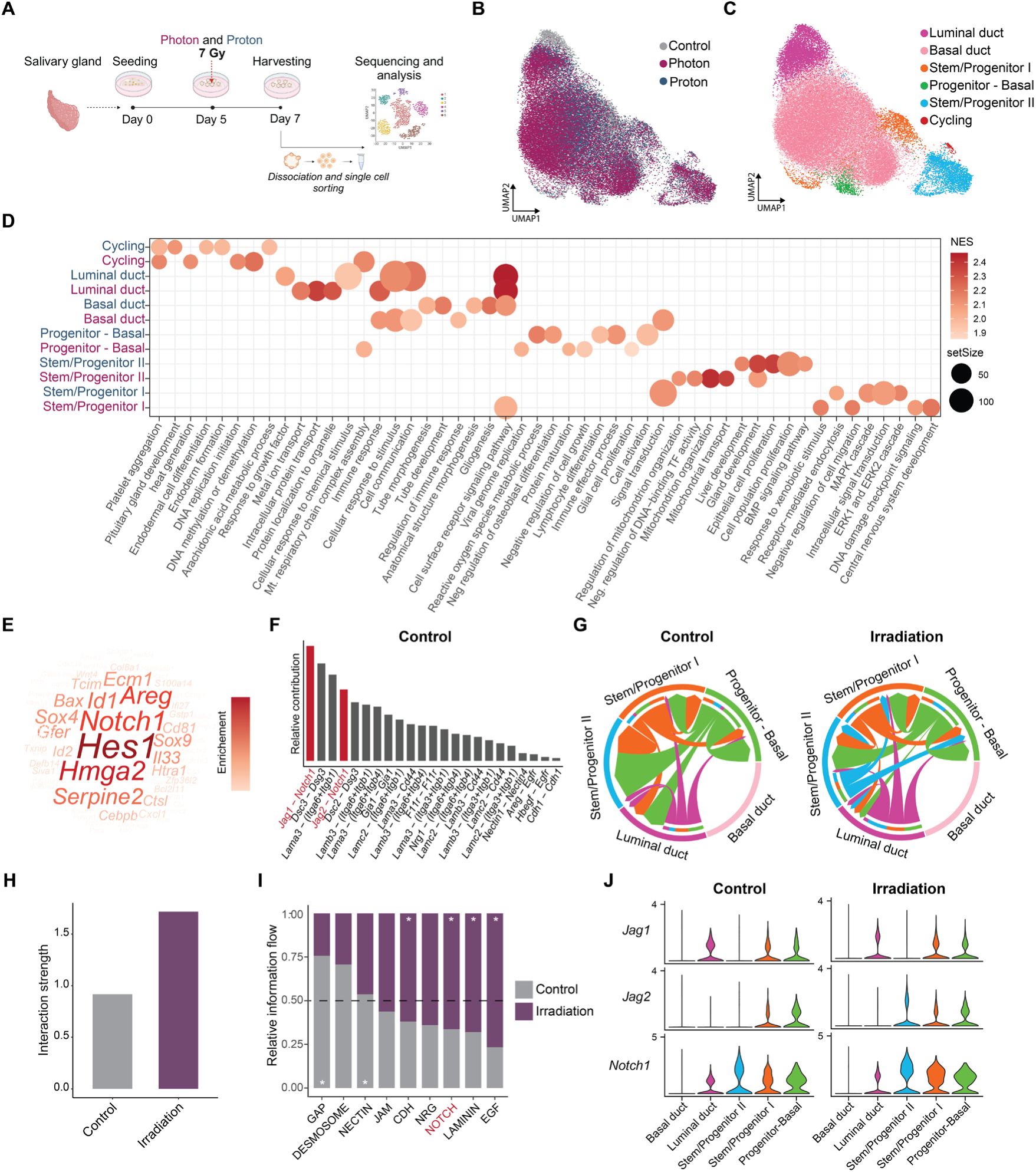
Surviving stem/progenitor cells show increased communication and Notch signaling after irradiation. (**A**) Schematic representation of the scRNA-seq experiment of photon and proton-irradiated mSGOs. (B) UMAP of control, photon and proton merged datasets showing the different conditions. (**C**) UMAP of control, photon and proton merged datasets showing the main cell populations. (**D**) Dot plot showing the top 5 significant (p.adj. < 0.05) biological processes in proton vs control (blue) and photon vs control (magenta) within each cell cluster. (**E**) Word cloud plot showing the most enriched genes in developmental, tissue morphogenesis and proliferation-related processes presented in Figure 3D. (**F**) Bar plot showing the significant ligand-receptor interactions in control mSGOs. Notch-related genes are shown in red. (**G**) Chord plot showing the interaction strength within each cell cluster in control and irradiated (photon + proton) mSGOs. (**H**) Bar plot showing the overall interaction strength in control and irradiated mSGOs. (**I**) Bar plot showing the most enriched signaling pathways shared between control and irradiated mSGOs. Paired Wilcoxon test was used for statistical analysis and * symbol indicates significant signaling pathways in control or irradiation. (**J**) Violin plots showing the expression of Notch signaling-related genes *Jag1*, *Jag2* and *Notch1* across the different populations in control and irradiated mSGOs.

To investigate potential signaling pathways important for stem/progenitor cell regulation in both control and irradiated samples, we performed a cell-cell interaction analysis using the 7-day control and irradiated mSGO datasets. Given the minimal differences observed between photon and proton irradiation, these samples were analyzed together and hereinafter referred to as irradiation group. Notch signaling, mediated by *Notch1-Jag1/Jag2* interaction, appeared as one of the most enriched pathways in control mSGOs (Figure 3F). Intriguingly, irradiation led to an overall heightened communication compared to control samples as shown by the increased communication patterns and strength (Figures 3G, 3H and S4D). Moreover, we observed a significantly higher enrichment of Notch signaling (Figure 3I), particularly within the stem/progenitor populations, which displayed increased outgoing and incoming signaling communication patterns (Figure S4E) and increased expression of Notch-related genes (Figure 3J, S4G). Of note, proton irradiation resulted in slightly higher expression of Notch-related genes and more pronounced communication patters compared to photon irradiation (Figures S4E, S4G). Taken together these results suggest a potential role for Notch signaling in regulating stem/progenitor cell activity after radiation damage.

### Salivary gland stem/progenitor cells rely on Notch signaling to maintain their self-renewal and migration capacity

Notch signaling plays a pivotal role in the regulation of stem cell activity and regeneration in several tissues^16^. Since Notch appeared as one of the most enriched signaling pathways in both control and irradiated mSGOs (Figures 3E, 3F), we investigated its role in maintaining the self-renewal capacity of salivary gland stem/progenitor cells *in vitro*. The self-renewal capacity of non-irradiated mSGOs, cultured in medium without Wnt and R-Spondin, factors that interact with Notch signaling and allow increased OFE^23^, was assessed upon treatment with the Notch γ-Secretase Inhibitor Dibenzazepine (DBZ) (Figures 4A). Notably, Notch inhibition (Figure S5A) impaired mSGO growth (Figure 4B) and significantly decreased the self-renewal capacity of salivary gland stem/progenitor cells, shown as a reduction of OFE and population doubling (PD) (Figures 4C). Next, to assess whether Notch inhibition influenced the differentiation capacity and stem cell exhaustion of mSGOs, untreated 7-day mSGOs were placed in differentiation media with or without DBZ and their ability to differentiate and self-renew were assessed (Figure 4D). In line with the previous findings, untreated organoids (-DBZ) showed a more spherical morphology (Figure 4E, left panel) and high levels of the basal/progenitor marker KRT14 (Figure S5B), indicative of a more premature and undifferentiated phenotype. In contrast, treatment with DBZ (+DBZ) (Figure 4E, right panel) or the Notch γ-Secretase Inhibitor DAPT (+DAPT) (Figure S5C) led to the formation of differentiated mini gland-like structures and the increased expression of the acinar marker AQP5 (Figure S5B). This aligned with previous work^28^, where Notch inhibition led to terminal differentiation and organoid maturation. Furthermore, mini glands treated with DBZ showed a much lower self-renewal capacity compared to untreated organoids (Figures 4F, 4G), confirming stem/progenitor cell exhaustion. To assess the impact of DBZ treatment on cell migration, we conducted a wound healing assay as previously described (Figures 2M, 2N). Notch inhibition reduced the motility of CD44-expressing cells (Figure 4H, S5D), with DBZ-treated samples showing a significantly lower capacity to fill the wound area compared to DMSO-treated samples (Figure 4I).Taken together, these data indicate that salivary gland stem/progenitor cells need Notch signaling to maintain their stemness, prevent the formation of differentiated structures, and sustain their motility in control conditions.

**Figure 4.**
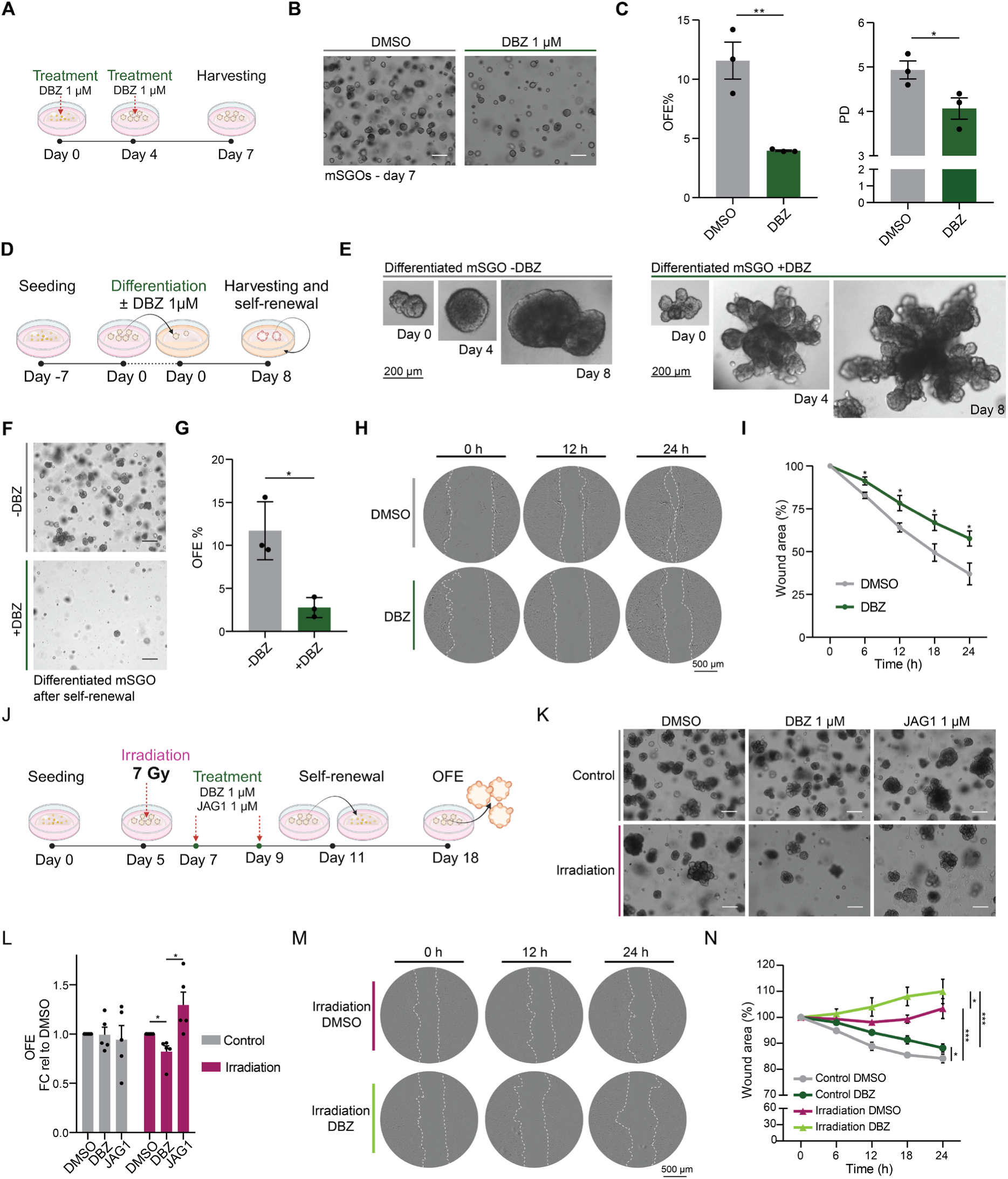
mSGOs rely on Notch signaling to maintain stemness and prevent premature differentiation. (**A**) Schematic representation of DBZ treatment in control mSGOs. (**B**) Representative images of 7-day mSGOs after treatment with DMSO or DBZ. Scale bar, 100 µm. (**C**) Organoid quantification of DMSO and DBZ-treated mSGOs shown as OFE% and population doubling (PD) (means ± s.e.m; n = 3 animals/condition). Two-sided unpaired *t*-test. * *p* < 0.05; ** *p* < 0.01. (**D**) Schematic representation of mSGO differentiation assay with or without DBZ treatment. (**E**) Representative images of differentiated mSGOs with or without DBZ treatment. Scale bar, 200 µm. (**F**) Representative images of differentiated mSGOs treated with or without DBZ after self-renewal. Scale bar, 100 µm. (**G**) Organoid quantification of differentiated mSGOs treated with or without DBZ after self-renewal shown as OFE% (means ± s.e.m; n = 3 animals/condition). Two-sided unpaired *t*-test. * *p* < 0.05. (**H**) Representative images of salivary gland cells after DMSO or DBZ treatment at 0, 12 and 24 h after wound generation. Dotted line shows wound borders. (**I**) Quantification of the wound area over time. Data is relative to 0 h (100%). One-way ANOVA, post-hoc Tukey’s test. * *p* < 0.05. (**J**) Schematic representation of DBZ and JAG1 treatment in control and photon-irradiated mSGOs. (**K**) Representative images of control and photon-irradiated mSGOs after treatment with DMSO, DBZ or JAG1. Scale bar, 100 µm. (**L**) Organoid quantification of control and photon-irradiated mSGOs after treatment with DMSO, DBZ or JAG1 shown as OFE. Data is shown as FC relative to DMSO (means ± s.e.m; n = 5 animals/condition). One-way ANOVA, post-hoc Tukey’s test. * *p* < 0.05. (**M**) Representative images of irradiated salivary gland cells after DMSO or DBZ treatment at 0, 12 and 24 h after wound generation. Dotted line shows wound borders. (**N**) Quantification of the wound area over time. Data is relative to 0 h (100%). One-way ANOVA, post-hoc Tukey’s test. * *p* < 0.05, *** *p* < 0.005.

After irradiation, Notch signaling was found to be highly upregulated (Figures 3I, S5E and S5F). To address if surviving stem/progenitor cells rely on Notch to maintain their self-renewal capacity after irradiation, mSGOs were irradiated at day 5 and treated with either the Notch inhibitor DBZ or the Notch activator JAG1 from day 7. Treated 11-day mSGOs were collected and self-renewal capacity was assessed (Figures 4J). Interestingly, DBZ treatment impaired the self-renewal capacity of irradiated mSGOs, shown as a reduction in OFE (Figures 4K, 4L, S5G and S5H). Additionally, Notch inhibition significantly impaired the migration abilities of irradiated salivary gland cells, which were already reduced compared to non-irradiated samples (Figures 4M, 4N), highlighting the critical role of Notch signaling in sustaining both stemness and migratory potential in irradiated salivary gland tissue. Notably, while JAG1 treatment significantly enhanced the OFE of photon-irradiated samples (Figures 4K, 4L), no changes were detected in proton-irradiated mSGOs (Figures S5G, S5H) that already showed significantly higher levels of Notch signaling at the single cell level (Figure S4G).

### Notch signaling maintains the stem cell potential of human salivary, thyroid and mammary gland-derived organoids

To extend our findings to other glandular tissues, we employed our previously described mouse thyroid gland organoid (mTGO) model^43,44^ and assessed the role of Notch signaling in regulating mTGO self-renewal capacity and differentiation *in vitro*. Similarly to mSGOs, DBZ treatment led to downregulation of Notch signaling (Figure S5I) and to a significant reduction of the self-renewal capacity of non-irradiated mTGOs (Figures 5A, 5B). Furthermore, DBZ-treated organoids showed increased lobular morphology (Figures 5A, 5C) and heightened expression of the thyroid differentiation markers *Slc5a5* (NIS) and *Tg* (Thyroglobulin) (Figure 5D), suggesting premature differentiation. To assess the differentiation capacity of mTGOs upon inhibition of Notch signaling, we placed untreated 7-day mTGOs in differentiation media^43^ with or without DBZ. Similarly to mSGOs (Figure 4E), DBZ-treated mTGOs showed a lobular phenotype (Figure 5E) accompanied by the upregulation of differentiation markers (Figure S5J).

**Figure 5.**
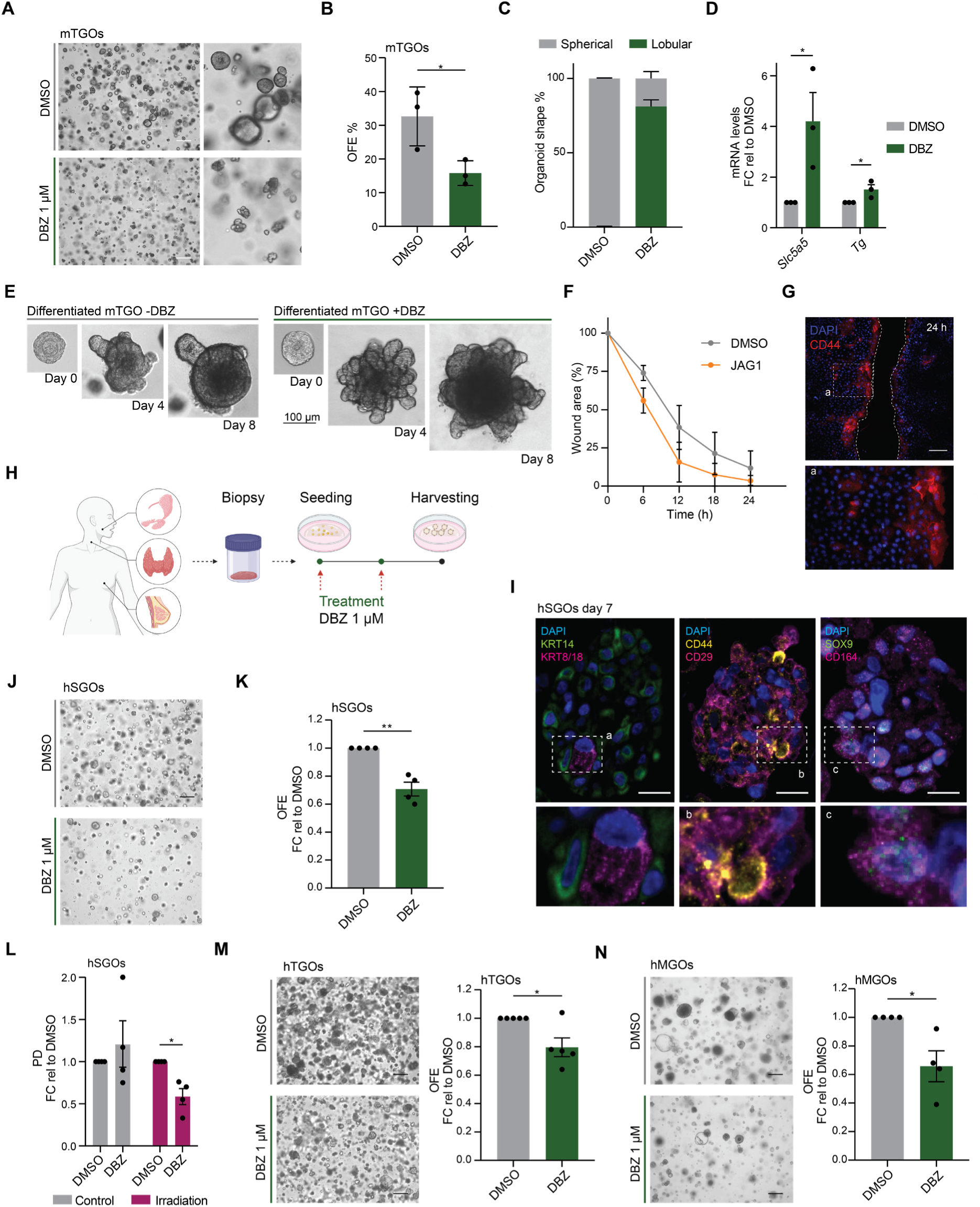
Notch signaling regulates stem cell self-renewal and differentiation of murine and patient-derived glandular organoids. (A) Representative images of 7-day-old mTGOs after treatment with DMSO or DBZ. Scale bar, 100 µm. (B) Organoid quantification of DMSO and DBZ-treated mTGOs shown as OFE% (means ± s.e.m; n = 3 animals/condition). Two-sided unpaired *t*-test. * *p* < 0.05. (**C**) Quantification of lobular and spherical mTGOs after DMSO and DBZ treatment. Data is shown as percentage (%) of lobular and spherical organoids/total organoid number. (**D**) rt-qPCR analysis of *Slc5a5* and *Tg* in mTGOs after DMSO and DBZ treatment. Data is shown as FC relative to DMSO (means ± s.e.m; n = 3 animals/condition). Two-sided unpaired *t*-test. * *p* < 0.05. (**E**) Representative images of differentiated mTGOs with or without DBZ treatment. Scale bar, 100 µm. (**F**) Quantification of the wound area over time. Data is relative to 0 h (100%). (**G**) Representative images of immunofluorescence staining of thyroid glands cells 24 h after wound generation and treatment with DMSO and JAG1, showing the expression of CD44. Scale bar, 100 µm. (**H**) Schematic representation of DBZ treatment in hSGOs, hTGOs and hMGOs. (**I**) Representative images of immunofluorescence staining of hSGOs showing the expression of KRT14, KRT8/18, CD29, CD44, SOX9 and CD164. Scale bar, 10 µm. (**J**) Representative images of hSGOs after DMSO and DBZ treatment. Scale bar, 100 µm. (**K**) Organoid quantification shown as OFE after DMSO and DBZ treatment. Data is shown as FC relative to DMSO (means ± s.e.m; n = 4 patients/condition). Two-sided unpaired *t*-test. ** *p* < 0.01. (**L**) Organoid quantification of control and photon-irradiated hSGOs after treatment with DMSO and DBZ shown as PD. Data is shown as FC relative to DMSO (means ± s.e.m; n = 4 patients/condition). Two-sided unpaired *t*-test. * *p* < 0.05. (**M**) Representative images of hTGOs (left) and organoid quantification shown as OFE (right) after DMSO and DBZ treatment. Data is shown as FC relative to DMSO (means ± s.e.m; n = 5 patients/condition). Scale bar, 100 µm. Two-sided unpaired *t*-test. * *p* < 0.05. (**N**) Representative images of hMGOs (left) and organoid quantification shown as OFE (right) after DMSO and DBZ treatment. Data is shown as FC relative to DMSO (means ± s.e.m; n = 4 patients/condition). Scale bar, 100 µm. Two-sided unpaired *t*-test. * *p* < 0.05.

Since our previous work identified CD44-expressing cells also in mTGOs^43^, we investigated whether these cells exhibited migratory properties similar to those in the salivary gland by performing a wound healing assay. Already 24 hours after wound generation, thyroid cells showed a pronounced migratory phenotype, which was further enhanced by Notch signaling activation with JAG1 (Figure 5F, S5K). Interestingly, as observed in salivary gland cells, CD44-expressing cells concentrated at the wound front (Figure 5G, S5L). Taken together, these findings emphasize the role of Notch signaling in mTGO self-renewal and differentiation and reveal a CD44-expressing cell population with migratory abilities.

A deeper comprehension of the signaling pathways governing patient-derived stem/progenitor cells can help improve the use and establishment of organoid models for preclinical studies and regenerative applications^6^. To translate our findings to patient-derived organoid models, we cultured organoids using biopsies obtained from the salivary glands of patients and assessed how Notch signaling influenced organoid growth and self-renewal capacity *in vitro* (Figure 5H). Similar to mSGOs, human submandibular salivary gland organoids (hSGOs) expressed basal and stem/progenitor cell markers (Figure 5I) and a significant decrease in self-renewal capacity following DBZ treatment (Figure 5J, 5K and S6B). To assess the regenerative capacity of human salivary gland stem/progenitor cells after irradiation, we treated irradiated hSGOs with DBZ and assessed their self-renewal capacity as previously described (Figure 4J). Notably, inhibition of Notch signaling after irradiation markedly impaired organoid growth, as evidenced by the significant decrease in PD after self-renewal (Figure 5L and S6A). These findings confirmed the conserved role of Notch signaling in regulating both murine and patient-derived SGOs in regenerative conditions. To extend these observations to other human glandular tissues, we generated organoids from patient biopsies of human thyroid (hTGOs) and mammary glands (hMGOs) (Figure 5H). Interestingly, all human glandular organoids exhibited an impairment in self-renewal and a significantly lower OFE upon DBZ treatment (Figures 5M, 5N and S6C, S6D), highlighting a conserved pre-regenerative role of Notch signaling across various patient-derived glandular organoid models.

## Discussion

Salivary gland stem/progenitor cells are a rare population of cells essential for normal tissue turnover in regenerative conditions. The establishment of SGOs has greatly advanced the characterization of these cells. Yet, the markers and the mechanisms defining them remain to be fully elucidated. This is the first study in which scRNA-seq profiling was used to characterize a mSGO model across different timepoints before and after irradiation. Bioinformatic and functional analyses identified *Sox9*, *Itgb1* and *Cd44* as novel markers of salivary gland stem/progenitor cells. Moreover, we demonstrated a conserved role of Notch signaling in regulating the regenerative capacities of both mouse and human stem/progenitor cells in organoids derived from exocrine and endocrine glandular tissues.

Compared to other more differentiated SGO models^28^, this model shows an enrichment of stem/progenitor cells, providing a valuable platform for the identification of markers of these rare populations, which are often difficult to detect and study in tissue samples. Indeed, scRNA-seq analysis revealed two distinct populations of salivary gland stem/progenitor cells, characterized by increased expression of several markers including *Sox9, Foxc1,* and *Cd24a* (stem/progenitor I), and *Itgb1, Cd44* and *Trp63* (stem/progenitor II). These genes have been previously identified as essential regulators of salivary gland development and branching morphogenesis processes^33,45–47^. Furthermore, *Sox9-expressing* cells have been shown to be involved in tissue regeneration in several organs following injury^48–50^, highlighting the regenerative stem cell features of this population. By mapping these genes into a scRNA-seq dataset of freshly isolated salivary gland tissue, we identified a small subpopulation of duct cells enriched with stem/progenitor markers. Although further studies are required to evaluate the presence of these cells *in vivo*, our results support the notion that a pool of active stem/progenitor cells might reside within the ductal compartment of adult salivary glands^1^. A more precise *in vivo* mapping of these cells could enhance our understanding of the regions to spare during radiotherapy of head and neck cancer, potentially improving both conventional and advanced radiotherapy techniques such as proton therapy^11^. *Itgb1* and *Cd44* marked a population of stem/progenitor cells with increased organoid formation and self-renewal capacity. Notably, this population showed elevated expression of acinar-like markers overtime, suggesting a potential role as pro-acinar precursor cells. Moreover, this cluster of cells also exhibited markers typical of mesenchymal-like cells, such as *Cd44, Alcam* (CD166) and *Nt5e* (CD73)^51^, and thus likely representing a plastic population of cells withtrans-differentiation and regeneration capacities^52^. Although these markers are shared across different cell types of various tissues, the wound healing assay clearly demonstrated the migratory potential of this population. Interestingly, the migratory potential of salivary gland stem cells may also explain the complete loss of homeostasis observed when only the top part of the salivary gland, which contains the tissue stem cells, is irradiated^11^. Together with their enhanced stemness and differentiation properties, this identifies a novel salivary gland cell population with both stem/progenitor and migratory features, which could potentially improve the efficiency of autologous transplantation and improve salivary gland regeneration following irradiation.

Notch signaling is a major pathway involved in stem/progenitor cell maintenance and differentiation across several tissues and organs^16^. While during development Notch signaling has been shown to be involved in cell fate decision and differentiation of multipotent precursors into salivary gland luminal and acinar cells^18^, our work reveals a role of Notch signaling in maintaining adult salivary gland stem/progenitor cell function in regenerative conditions. Our findings, which extend to various murine and human-derived organoids, support previous studies^28,53–55^ demonstrating that the inactivation of the Notch pathway induces terminal differentiation of stem/progenitor cells in different tissues. Furthermore, in line with prior research establishing a critical role of Notch signaling in regulating thyroid and mammary gland function^56,57^, we demonstrated its significant role in maintaining the self-renewal potential of both TGOs and MGOs. Additionally, we revealed a novel role of Notch signaling in regulating the regenerative capacities of both murine and human salivary gland stem/progenitor cells following irradiation, which could improve future radiotherapy applications. Intriguingly, cell-cell interaction analysis uncovered a higher enrichment of Notch signaling after proton irradiation, suggesting enhanced stem/progenitor cell maintenance, and a potential advantage of proton therapy over conventional photon-based radiotherapy. Previous studies have shown a link between Notch signaling and radiation-induced DNA damage^58^, which might influence later stem/progenitor cell responses. Notably, in our previous work^15^, we demonstrated that *Sox9-* expressing cells exhibit a higher interferon beta (IFNB) response and increased compensatory proliferation following proton irradiation at later timepoints. Although further studies are required to establish a connection between Notch signaling, IFNB signaling and stem/progenitor cell activity, we speculate that early activation of Notch signaling could influence radiation-induced immunogenic responses of normal tissue stem/progenitor cells^59^, thereby influencing their late response and self-renewal capacities. In summary, our work highlights the complexity of mSGOs and reveals the presence of novel stem/progenitor cell populations within these organoids. Additionally, it underscores the essential role of Notch signaling in regulating the regenerative capacity of murine and patient-derived stem/progenitor cells in different glandular mouse and human organoid models.

### Limitations of the study

While our work has identified novel putative salivary gland stem/progenitor cells, further studies are required to validate the presence and function of these cell populations *in vivo*. Moreover, although our findings support previous data on the role of Notch signaling in regulating adult stem/progenitor cells, translating these results *in vivo* is essential to understand how a more complex microenvironment and neighboring cells influence this signaling pathway under stress conditions.

To exclude dead cells from the scRNA-seq analysis of irradiated mSGOs, FACS was conducted immediately prior to sequencing. The FACS procedure might have influence cell integrity and transcriptional changes. However, through accurate performance and handling the cells on ice, we ensured reproducibility and minimize cellular stress across all conditions.

Considering the strong link between Notch signaling and hypoxia^60^, it is essential to take into account the lower oxygen levels of tissue culture conditions compared to *in vivo* settings, as they could potentially influence the regulation of Notch signaling. Additionally, due to known interaction between Wnt and Notch signaling pathways^61^, the presence of Wnt in the medium of irradiated organoids might have influenced salivary gland growth even after Notch inhibition.

Due to technical restrictions, only female mice were used to generate organoids, whereas both male and female patients were included for human-derived organoids. Potential sex differences in stem/progenitor cell activity remain to be evaluated.

## Acknowledgments

We would like to thank the surgical teams of the University Medical Center Groningen, Medical Center Leeuwarden and Martini Ziekenhuis for providing donor biopsies. We also thank S. Both, J. Free and B.N. Jones for the technical support at the PARTREC and GPTC; B. Eggen, N. Brouwer and E. Gerrits for support in the scRNA-seq library preparation and analysis; the members of the ERIBA Research Sequencing Facility (UMCG) for the sequencing runs; T. Bijma and J. Teunis from the Flow Cytometry Unit (UMCG) for the sorting. Illustrations were created with Biorender.com. This work was supported by the Dutch Cancer Society KWF Grant nr 12092 and IBA Grant nr PPP-2021-27.

## Authors contribution

D.C, L.B and R.P.C conceived and designed the study with advice and input from S.P. and A.S.G.; D.C., L.K., Y.W. and I.R.H. performed the sequencing experiments and analyses with advice and input from A.S.G.; D.C. and M.J.v.G. performed the irradiations; D.C., R.M., J.P.G., P.S.M. and A.L.J.d.B performed the mouse, thyroid and mammary gland experiments; D.C., L.B. and R.P.C. wrote the manuscript. All authors approved the manuscript.

## Material and methods

Details of all reagents and antibodies used in this study are mentioned in Table S1.

### Mice

Submandibular salivary glands and thyroid glands were collected from 8-to 12-week-old female C57BL/6 mice (Envigo, the Netherlands). Mice were housed under standard conditions with diet and water *ad libitum*. Animal work was approved by the Central Committee of Animal Experimentation of the Dutch government and the Institute Animal Welfare Body of the University Medical Center Groningen (UMCG) [animal welfare body (IVD) protocol number 184824-01-001].

### Patient biopsies

Human non-malignant submandibular gland, thyroid gland and mammary tissues were obtained from donors after informed consent and Institutional Review Board (IRB) approval during scheduled surgery for the removal of squamous cell carcinoma of the oral cavity (for the submandibular gland), papillary thyroid carcinoma (for the thyroid gland) and invasive ductal carcinoma (for the mammary gland) at the University Medical Center Groningen (UMCG), Medical Centre Leeuwarden and Martini Ziekenhuis.

### Organoid cultures

*Mouse salivary gland organoids:* mouse submandibular salivary glands were first mechanically and enzymatically digested in digestion buffer [collagenase type II (0.63 mg/mL), hyaluronidase (0.5 mg/mL) and CaCl_2_ (6.25 mM) in HBSS/BSA 1%] and then cultured in minimal medium (MM) consisting of DMEM/F12, penicillin-streptomycin (Pen/Strep) antibiotics, glutamax (2 mM), EGF (20 ng/ml), FGF2 (20 ng/ml), N2 (1x), insulin (10 µg/ml) and dexamethasone (1 µM). After 3 days in culture, mouse salivary spheres were dissociated into single cells using 0.05% trypsin EDTA and counted. 10.000 mouse cells were then plated in 75 µL gel/well [25 µL cell suspension + 50 µL of Matrigel] in a 12-well tissue culture plate. After solidification of the gels, 1 mL of WRY medium [MM, Y27632 (10 µM), 10% R-spondin1– conditioned medium, and 50% Wnt3a-conditioned medium] was added to each well. The plates were then placed at 37°C and 5% CO_2_ until the day of the experiment.

*Human salivary gland organoids:* human submandibular salivary gland biopsies were mechanically and enzymatically digested in digestion buffer and counted. 20.000 cells were then plated in 75 µL gel/well [25 µL cell suspension + 50 µL of Matrigel] in a 12-well tissue culture plate. After solidification of the gels, 1 mL of WRYTN medium [MM medium, Y27632, Noggin (50 ng/mL), A8301 (1 µM), 10% R-spondin1–conditioned medium, and 50% Wnt3a-conditioned medium] was added to each well of human cells. The plates were then placed at 37°C and 5% CO_2_ until the day of the experiment.

*Mouse thyroid gland organoids:* as previously described^43^, following mechanical and enzymatic digestion mouse samples were cultured in thyroid gland medium (TGM) consisting of DMEM/F12, Pen/Strep antibiotics, glutamax, EGF, FGF2, and 0.5% B27 supplement. One day after isolation, mouse thyroid spheres were dissociated into single cells using 0.05% trypsin EDTA and counted. 10.000 mouse cells were then placed in 75 µL gel/well [25 µL cell suspension + 50 µL of Matrigel] in a 12-well tissue culture plate and 1 mL of TGM supplemented with Y27632 was added to each well. The plates were placed at 37°C and 5% CO_2_ until the day of the experiment.

*Human thyroid gland organoids:* as previously described^43^, following mechanical and enzymatic digestion, human thyroid gland cells were counted and 25.000 cells were placed in 75 µL gel/well [25 µL cell suspension + 50 µL of Matrigel] in a 12-well tissue culture plate. 1 mL of human TGM (hTGM) consisting of DMEM/F12, Pen/Strep antibiotics, glutamax, R-Spondin-1 human (100 ng/mL), 10% UltraGRO-PURE GI Cell Culture Supplement Xeno-free, EGF (40 ng/mL), FGF2, 0.5% CTS B27 supplement XenoFree, Y27632, A8301 (5 µM), nicotinamide (10 mM) and 1% Heparin Sodium Salt Solution was then added to each well. The plates were then placed at 37°C and 5% CO_2_ until the day of the experiment.

*Mammary gland organoids:* mammary gland biopsies were first mechanically and enzymatically digested in digestion buffer consisting of mammary gland medium (MGM) [Advanced DMEM, Pen/Strep antibiotics, glutamax, 20% Wnt3a-conditioned medium, 10% R-spondin conditioned medium, 1x B27 supplement, nicotinamide, Noggin (25 ng/mL), Y27632 (5 µM), N-acetylcysteine (1,25 mM), Primocin (100 µg/mL), Hydrocortisone (500 ng/mL), β-estradiol (100 nM), Forskolin (10 µM), Heregulin β1 (5 nM), EGF (5 ng/mL), SB202190 (1 µM), FGF7 (5 ng/mL), FGF10 (20 ng/mL), A8301 (0.5 µM) and 1% Heparin Sodium Salt Solution] and Collagenase II (1 mg/mL). Human mammary gland cells were then counted and 10.000 cells were placed in 80 µL gel/well [20 µL cell suspension + 60 µL growth factor reduced BME extract] in a 12-well tissue culture plate. After solidification of the gels 1 mL of MGM was added to each well. The plates were placed at 37°C and 5% CO_2_ until the day of the experiment.

### Self-renewal assay

To assess the self-renewal capacity of the stem/progenitor cells within the mouse and human organoid models, organoids were collected and dissociated into single cells at the proper time point using 0.05% trypsin-EDTA. As previously described, single cells were then placed in Matrigel (SGOs and TGOs) or BME (MGOs) in a 12-well tissue culture plate with 1 mL of full medium. Plates were placed at 37°C and 5% CO_2_. Between 7 and 14 days later (based on the organoid model), organoids were collected and dissociated into single cells. Organoid and single cell numbers were noted and used to calculate the organoid formation efficiency (OFE) and population doubling (PD) as follows:

OFE%= (total number of organoids harvested / number of seeded cells) x 100

PD = ln (total number of harvested cells / number of seeded cells) / ln2

### Organoid differentiation

To assess the differentiation capacity of both mouse and thyroid gland organoids, 7-day organoids were carefully harvested and counted. Approximately 30 to 50 organoids were then placed in a 75 µL gel/well [25 µL cell suspension + 50 µL of Matrigel] in a 96-well tissue culture plate previously coated with a layer of Matrigel diluted 1:1 (20 µL DMEM/F12 + 20 µL Matrigel). After solidification of the gels, 150 µL of salivary gland differentiation media [MM, 10% FBS, HGF (50 ng/mL), ±DBZ (1 µM)] or 150 µL of thyroid gland differentiation media [TGM, 10% FBS, Heparin Sodium Salt Solution (100 ng/mL), IGF1 (50 ng/ml), FGF10 (100 ng/mL), insulin (5 µg/ml), ITS (5 µg/ml), Dexamethasone (50 nM), bTSH (100 mU/mL), ±DBZ (1 µM)] were added to each well and refreshed every 2-3 days. The plates were then placed at 37°C and 5% CO_2_. After 8 days in culture, the organoids were collected and processed for rt-qPCR or fixed and embedded in paraffin for immunofluorescence staining.

### Treatments

5-day salivary gland organoids were irradiated either with 7 Gy photons or protons. Photon irradiation was performed using a Cesium-137 source with a dose rate of 0.59 Gy/min at the department of Biomedical Sciences of the UMCG. Proton irradiation was performed with a 150 MeV proton beam at the Particle Therapy Research Center (PARTREC) accelerator facility of the UMCG or at the UMCG Proton Therapy Center (GPTC). Samples were placed in the plateau region of a 150 MeV Bragg curve.

To assess the role of Notch signaling in irradiated samples Dibenzoazepine (DBZ), JAG1 or equivalent volume of solvent (DMSO) were added to salivary gland organoids at a final concentration of 1 µM. To evaluate the role of Notch signaling in control condition, organoids from all glands were cultured without Wnt3a-conditioned medium [Enriched Medium (EM)]. 1 µM of DBZ was added to the medium from day 0 and refreshed every 4 days until the end of the experiment. mSGOs were also treated with 1 µM of DAPT.

### Fluorescence activated cell-sorting

For the sorting of stem/progenitor cells, mouse salivary gland organoids were harvested and dissociated into single cells using 0.05% trypsin EDTA. Cells were resuspended in 0.2% PBS/BSA and incubated with the proper conjugated primary antibody (Table S1) for 30 min at 4°C. After washing, cells were resuspended in 0.2% PBS/BSA containing propidium iodide (PI; 1 mg/mL) and sorted with the Moflo Astrios sorter at the Flow Cytometry Unit (FCU) at the UMCG. Sorted cells were washed and placed in a 75 µL gel/well [25 µL cell suspension + 50 µL of Matrigel] at a concentration of 10.000 cells/gel in a 12-well tissue culture plate. 1 mL of WRY media was added to each well and the plates were then placed at 37°C and 5% CO_2_. Organoids were harvested and counted at day 7 for OFE calculation. Flow cytometry analysis was performed using the FlowJo software (v10.10.0). The list of antibodies and dilutions are specified in Table S1.

### Wound healing assay

Control and irradiated salivary gland and thyroid gland organoids were collected and dissociated into single cells using 0.05% trypsin-EDTA. After cell counting, 90,000 cells/well were re-seeded in a 24-well plate pre-coated with Matrigel. WRY medium (for salivary gland cells) or TGM medium (for thyroid gland cells) was added, and cells were incubated at 37°C with 5% CO₂. Once cells reached confluency, they were starved by replacing the full medium with DMEM/F12 without growth factors for at least 3 hours. A wound was then created in each well, and fresh DMEM/F12 containing either DBZ (1 µM), JAG1 (1 µM), or DMSO was added. Plates were then placed in an IncuCyte S3 Live-Cell Analysis System for imaging. After 24 hours, cells were fixed with 4% formaldehyde at room temperature for 10 minutes and then permeabilized using 0,2% Triton for 4 minutes. Primary antibody diluted in blocking solution (2% BSA) was then added to each well and incubated at room temperature for 1 hour. After washing, appropriate secondary antibody diluted in blocking solution (2% BSA) was added to each well at room temperature for 1 hour and with DAPI at room temperature for 10 minutes for nuclear staining. Images were then acquired using a Leica DMI 8 Inverted microscope. The list of antibodies and dilutions is specified in Table S1. Images were processed and analyzed using ImageJ (v1.52).

### Immunofluorescence staining

For the staining of organoid sections, salivary and thyroid gland organoids were collected, fixed with 4% formaldehyde for 15 minutes and transferred into a drop of HistoGel embedding medium. HistoGels were then dehydrated, embedded in paraffin and cut into 4 µm thick sections. For the staining, sections were dewaxed and cooked with Tris-EDTA (pH 9) antigen retrieval buffer. After permeabilization and blocking [4% donkey serum, 1% BSA, 0.01% Triton in PBS 1x], sections were incubated with the proper primary antibody at 4°C overnight. On the next day, sections were incubated with the secondary antibody at room temperature for 1 hour and with DAPI at room temperature for 10 minutes for nuclear staining. Upon mounting, images were acquired with a Leica DM6 microscope. The list of antibodies and dilutions are specified in Table S1. Images were processed using ImageJ (v1.52).

### Quantitative real-time qPCR

Isolation of total RNA from salivary and thyroid gland organoids was performed using the RNeasy Mini Kit according to the manufacturer’s protocol. RNA was retro transcribed by using 1 µL dNTP mix (10 mM), 1 µL random primers (100 ng), 4 µL 5x First-strand Buffer, 2 µL DTT (0.1 M), 1 µL RNase OUTTM (40 units/µL), and 1 µL M-MLV RT (200 units). To assess the expression of the genes of interest, specific primers were used together with iQ SYBR Green Supermix. All reactions were run in triplicate on a Bio-Rad Real-Time PCR System. The list of primers is specified in Table S2. The *Ywhaz* gene was used as internal control.

### Single-cell RNA sequencing library preparation

The preparation of the single-cell RNA sequencing library was performed as previously described^15^. In brief, upon harvesting and dissociation of the salivary gland organoids, 45.000 propidium iodide negative cells/sample were sorted using the Moflo Astrios sorter at the FCU at the UMCG. Samples were then loaded on the Chromium Next GEM Chip G and ran in the Chromium Controller (10X Genomics). Library construction was performed following the 10X Genomics protocol and library quality control and concentration measurement were performed using the Qubit and Tapestation machines at the Research Sequencing Facility of ERIBA at the UMCG. For the sequencing, libraries were equimolarly pooled and 1.8 pM of the pool with 5% PhiX were loaded on a NextSeq 500 (Illumina) for a 75 bp paired-end sequencing run at the Research Sequencing Facility of ERIBA (UMCG).

### Single-cell RNA sequencing analysis

Reads without cell barcodes or UMIs were excluded and remaining raw reads were aligned to the mouse genome GRCm39. Following demultiplexing and cell filtering, the remaining cells were used for downstream analysis. Single-cell RNA sequencing analysis was done on R (v4.3.2) using the Seurat package (v5.0.3)^62^. Cells were excluded based on their mitochondria counts (more than 5%) and feature counts (less than 2500). The principal components (PC) to use for UMAP dimensionality reduction for each sample were calculated using the Elbow plots and Jack Straw. Doublets and clusters with less than 50 cells with no significant markers were excluded from the analysis and cluster-markers were identified using the FindAllMarkers function with the statistical test MAST. The CellChat package (v2.1.2)^63^ was used for cell-cell interaction analysis. Harmony^64^ (v.1.2.0) was used for the integration of 7-day and 11-day mSGOs. Pseudotime and trajectory analyses were performed on Python (v3.8.10) using the package pyVIA (v0.1.96)^65^.

### Bulk RNA-sequencing analysis

Previously deposited RNA-seq data was used for the analysis (GSE******). Analysis was performed as previously described^15^. In brief, for the alignment of the fastq files the mouse genome GRCm39 was used. R (v4.3.2) and RStudio were used for the downstream analysis. For the differential expression analysis, low expressed genes (total count <1 in less than 2 samples) were excluded. The R package edgeR^66^ was used for normalization and identification of differential expressed genes (LogFC > 0.6, FDR < 0.05).

### ATAC-sequencing library preparation and analysis

ATAC-sequencing (ATAC-seq) library preparation was performed as previously described^67^. In brief, mSGOs derived from 4 different mice were harvested. ATAC-seq libraries were then prepared using Nextera DNA Sample Preparation Kit (Illumina, FC-121-1030) following the manufacturer’s protocol. Following PCR amplification, DNA was purified using the Qiagen MinElute kit and run on a 2% E-gel agarose gel (Thermo Fished Scientific, G521802). The concentration of the libraries was determined using a Qubit (ThermoFisher Scientific) and 2100 Bioanalyzer (Agilent). 1,6 pM of libraries with 5% PhiX was then loaded on a NextSeq 500 (Illumina) for a 75 bp paired-end sequencing run at the Research Sequencing Facility of ERIBA (UMCG).

For the analysis, alignment and peak calling were performed using the ENCODE ATAC-seq pipeline. All organoid samples were used to generate a consensus peak file (>41K peaks). HOMER^68^ function findMotifsGenome was used to perform motif enrichment analysis with default settings. Transcription factors associated to each motif were integrated with the single-cell RNA sequencing dataset and their average expression within each cell cluster was calculated.

### Statistical analysis

The number of biological replicated and the details of the statistical tests used in each experiment are stated in main and supplementary figure legends. Data is shown as mean ± s.e.m., the Shapiro-Wilk test was used to test normality of distribution of raw data. Two-sided unpaired *t*-test was used for two group comparison while one-way ANOVA and two-way ANOVA with post-hoc Tukey’s test were used for multiple (> 2) group comparison. Sample sizes were estimated empirically and p-values ≤ 0.05 were considered statistically significant. Statistical analyses were performed using the GraphPad Prism software (v8.0.1).

## Data availability

The sequencing data have been deposited in the Gene Expression Omnibus (GEO) repository; GSE numbers are available for review purposes and will be made fully available upon acceptance of the manuscript. All other data are stored at the department of Biomedical Sciences, UMCG and are available from the corresponding authors upon request.

The details of all antibodies and reagents used in this study are listed in Table S1.

## Supplementary material

Figures S1-S6 Tables S1, S2

