## Supplementary Figures and Tables for "Notch Signaling Drives Pro-Regenerative and Migratory Traits in Glandular Stem/Progenitor cells"

Davide Cinat *et al.*

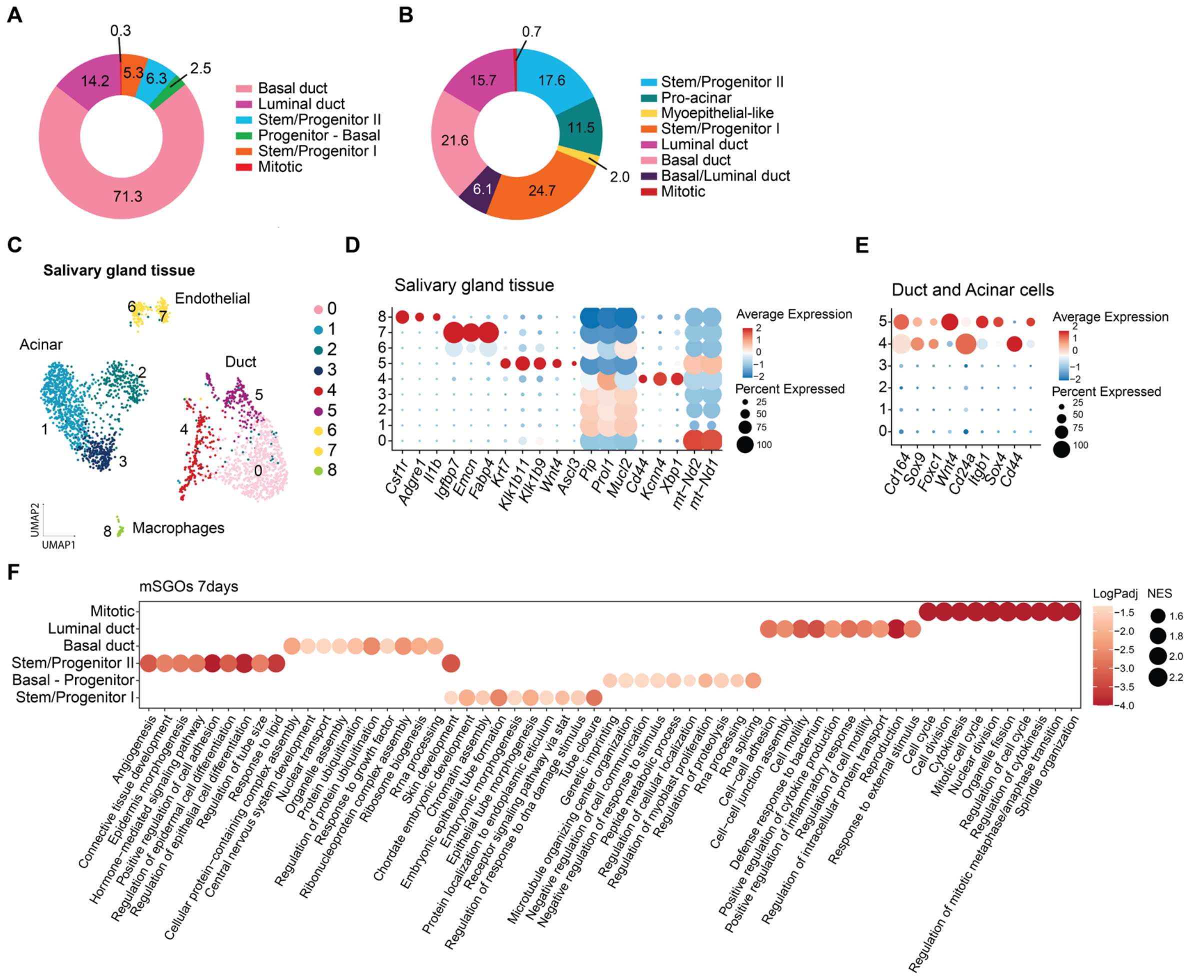

**Figure S1. scRNAseq analysis of mSGOs and salivary gland tissue**

(**A**) Cell proportion of clusters in 7-day mSGOs, UMAP is shown in Figure 1D. (**B**) Cell proportion of clusters in 11-day mSGOs, UMAP is shown in Figure 1E. (**C**) UMAP of the salivary gland tissue dataset showing the main cell populations. (**D**) Dot plot showing cell type marker genes of the salivary gland tissue dataset. (**E**) Dot plot showing stem/progenitor cell markers extrapolated from 7-day mSGOs in ductal and acinar subpopulations. (**F**) Dot plot showing the top 10 upregulated biological processes in each population of 7-day mSGOs.

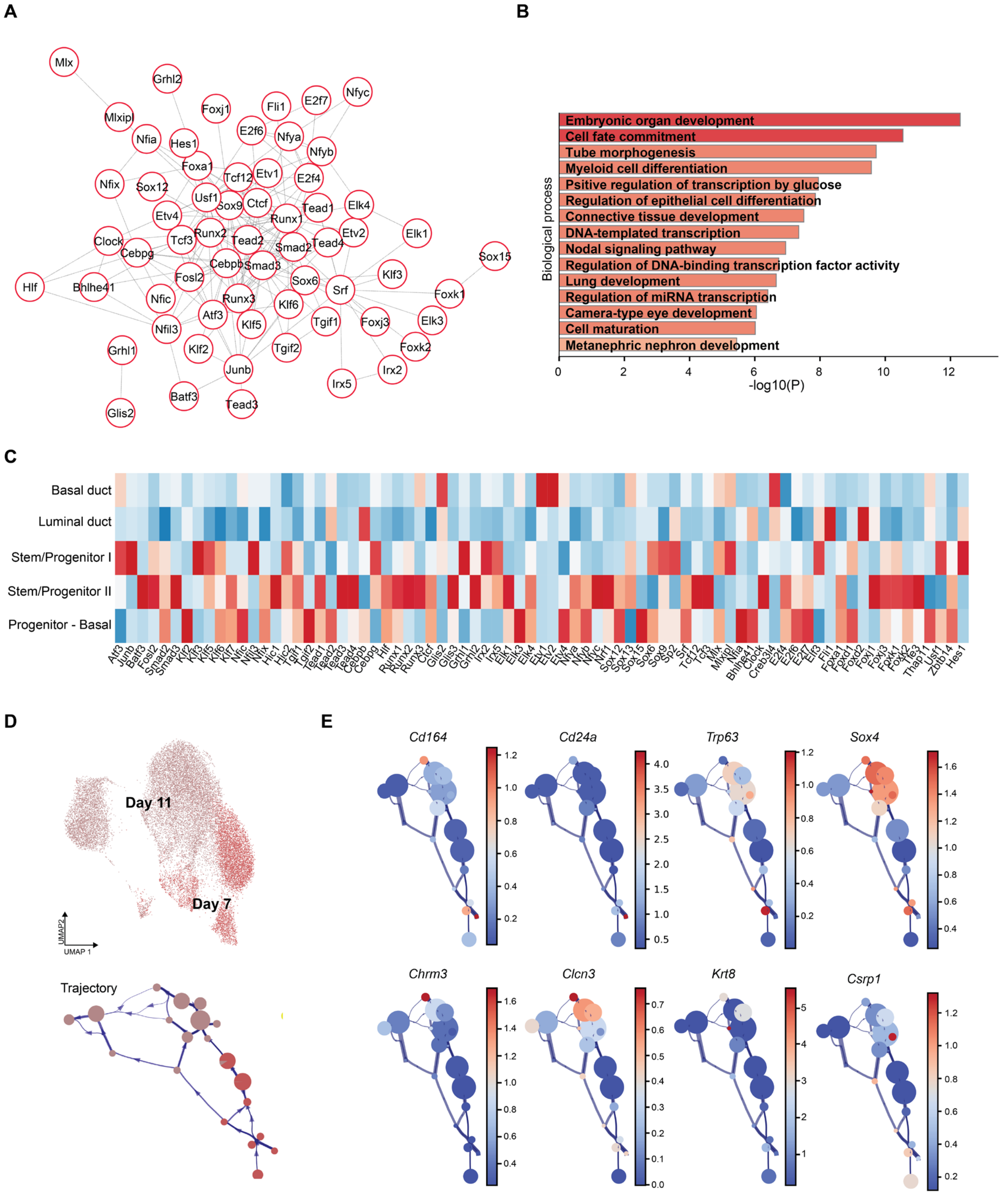


**Figure S2. 7-day mSGO populations show stem/progenitor cell features**

(**A**) Network analysis showing transcription factors related to significant ATAC-seq motifs. (**B**) Top 15 biological processes related to ATAC-seq motif transcription factors. (**C**) Heatmap showing the average expression of transcription factors related to significant ATAC-seq motifs across 7-day mSGO populations. (**D**) UMAP of the 7-day and 11-day mSGO-integrated dataset (same as in Figure 2B) and trajectory plot showing cluster distribution. (**E**) Trajectory plot of the 7-day and 11-day mSGO-integrated dataset showing the expression of stem/progenitor-related genes (*Cd164, Cd24a, Trp63, Sox4*) and acinar/duct-related genes (*Chrm3, Clnc3, Krt8, Csrp1*).

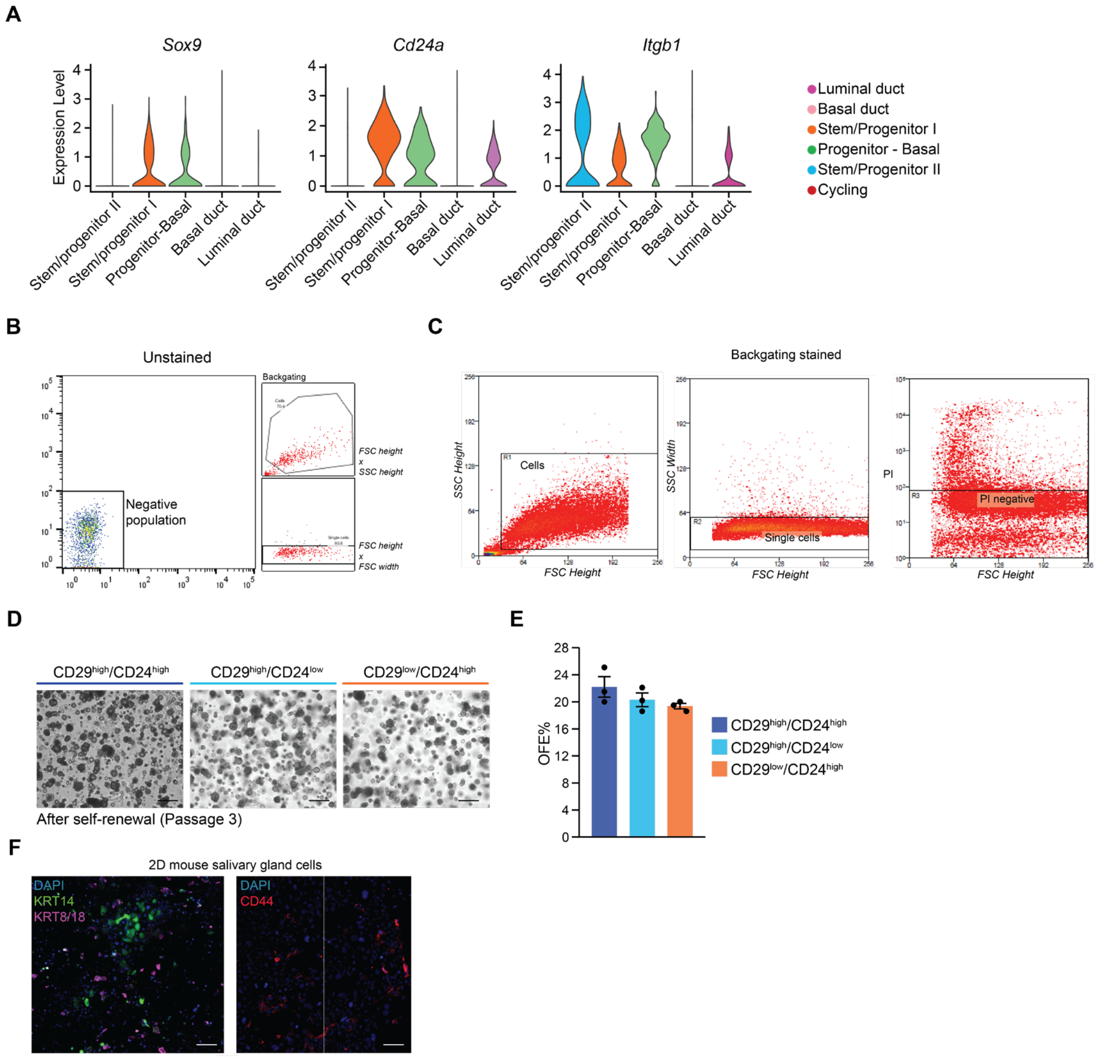


### Figure S3. FACS of 7-day mSGOs and 2D salivary gland cells

(**A**) Violin plot showing expression of *Sox9, Cd24a* (CD24) and *Itgb1* (CD29) across the 7-day mSGO populations. (**B**) Representative FACS plot of an unstained sample using the same gating strategy shown in Figure 2G. (**C**) Backgating of FACS plot shown in Figure 2G. (**D**) Representative images of sorted cells after self-renewal (P3). Scale bar, 100 µm. (**E**) Organoid quantification of sorted cells after self-renewal (P3) shown as organoid formation efficiency (OFE%) (means ± s.e.m; n = 3 animals/condition). (**F**) Representative images of immunofluorescence staining of 2D salivary gland cells showing the expression of KRT14, KRT8/18 and CD44. Scale bar, 100 µm.

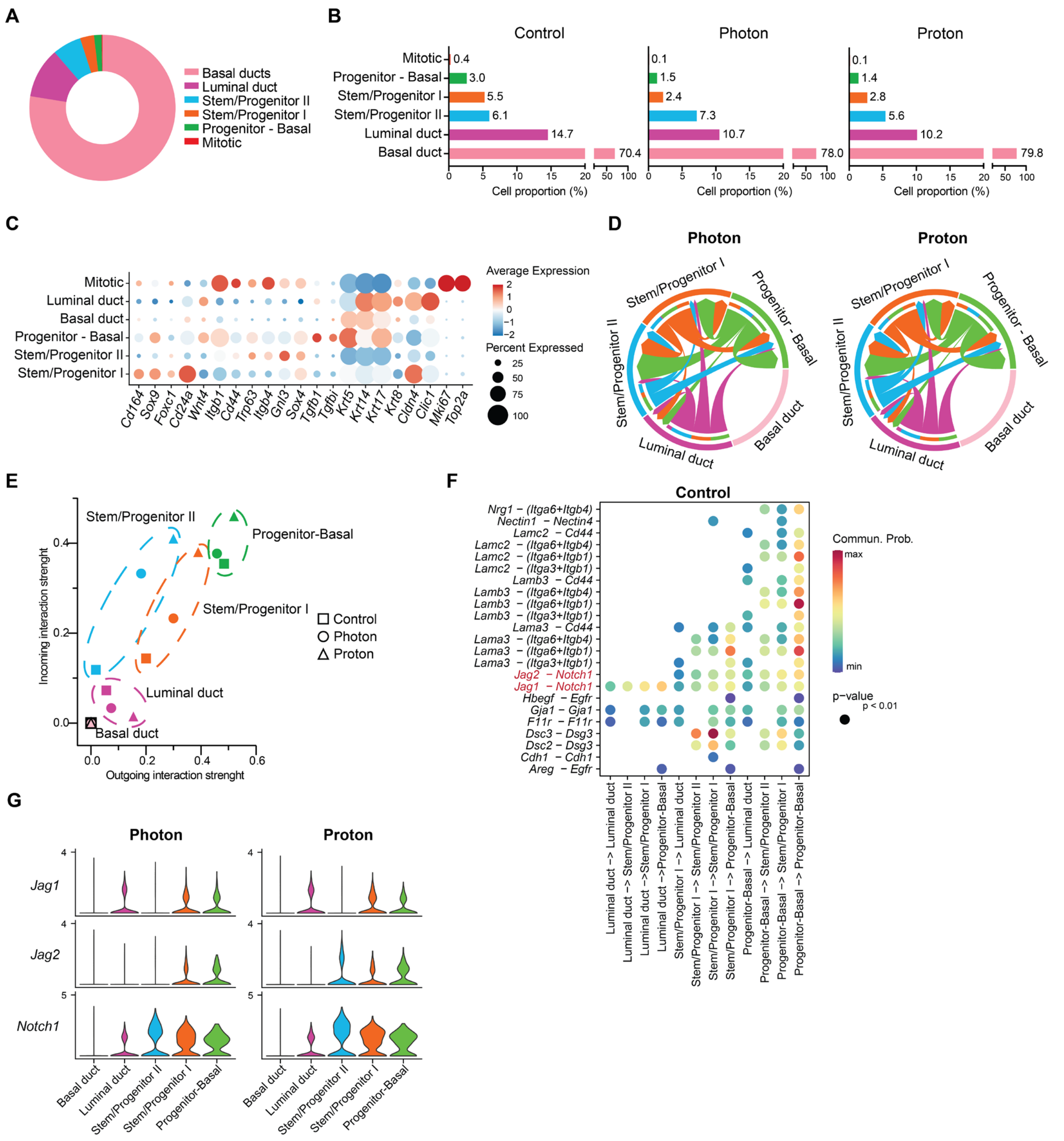


**Figure S4. scRNAseq analysis of photon and proton-irradiated mSGOs**

(**A**) Cell proportion of clusters in the merged control, photon and proton dataset, UMAP is shown in Figure 3C. (**B**) Cell proportion of clusters in control, photon and proton irradiated-mSGOs, UMAP is shown in Figure 3C. (**C**) Dot plot showing cell marker genes of the merged control, photon and proton dataset, UMAP is shown in Figure 3C. (**D**) Chord plot showing interaction strength in photon and proton-irradiated mSGOs. (**E**) PCA plot showing outgoing and incoming interaction strengths. (**F**) Bubble plot showing significant ligand-receptor interactions across the clusters of control mSGOs. (**G**) Violin plot showing the expression of Notch-related genes *Notch1, Jag1* and *Jag2* in photon and proton-irradiated mSGOs.

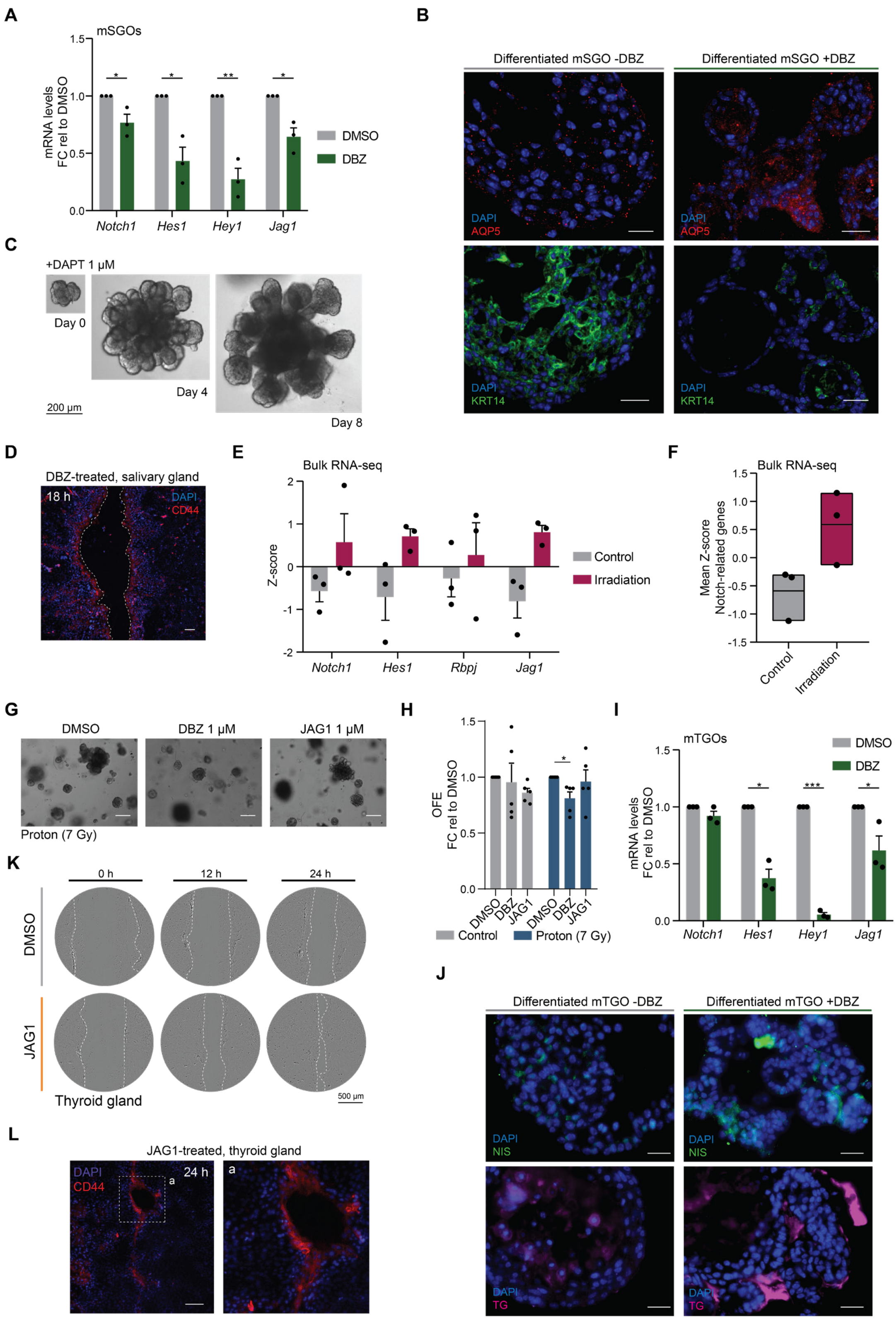


### Figure S5. Notch inhibition in mSGOs and mTGOs

(**A**) rt-qPCR analysis of Notch-related genes in mSGOs after DMSO and DBZ treatment. Data is shown as FC relative to DMSO (means ± s.e.m; n = 3 animals/condition). Two-sided unpaired *t*-test. * $p < 0.05$; ** $p < 0.01$. (**B**) Representative images of immunofluorescence staining of differentiated mSGOs with or without DBZ treatment showing the expression of KRT14 and AQP5. Scale bar, 10 µm. (**C**) Representative image of differentiated mSGOs after treatment with DAPT. Scale bar, 200 µm. (**D**) Representative images of immunofluorescence staining of salivary glands cells 18 h after wound generation and DBZ treatment showing the expression of CD44. Scale bar, 100 µm. (**E**) Gene expression of Notch-related genes extrapolated from the bulk RNA-seq analysis of control and photon-irradiated mSGOs. Data is shown as Z-score (n = 3 animals/condition). (**F**) Mean Z-score of the Notch-related genes shown in Figure S5E. (**G**) Representative images of proton-irradiated mSGOs after treatment with DMSO, DBZ and JAG1. Scale bar, 100 µm. (**H**) Organoid quantification of control and proton-irradiated mSGOs after treatment with DMSO, DBZ or JAG1 shown as OFE. Data is shown as FC relative to DMSO (means ± s.e.m; n = 5 animals/condition). One-way ANOVA, post-hoc Tukey`s test. * $p < 0.05$. (**I**) rt-qPCR analysis of Notch-related genes in mTGOs after DMSO and DBZ treatment. Data is shown as FC relative to DMSO (means ± s.e.m; n = 3 animals/condition). Two-sided unpaired *t*-test. * $p < 0.05$; *** $p < 0.005$. (**J**) Representative images of immunofluorescence staining of differentiated mTGOs with or without DBZ treatment showing the expression of NIS and TG. Scale bar, 20 µm. (**K**) Representative images of thyroid gland cells after DMSO or JAG1 treatment at 0, 12 and 24 h after wound generation. Dotted line shows wound borders. (**L**) Representative images of immunofluorescence staining of thyroid glands cells 24h after wound generation and JAG1 treatment showing the expression of CD44. Scale bar, 100 µm.

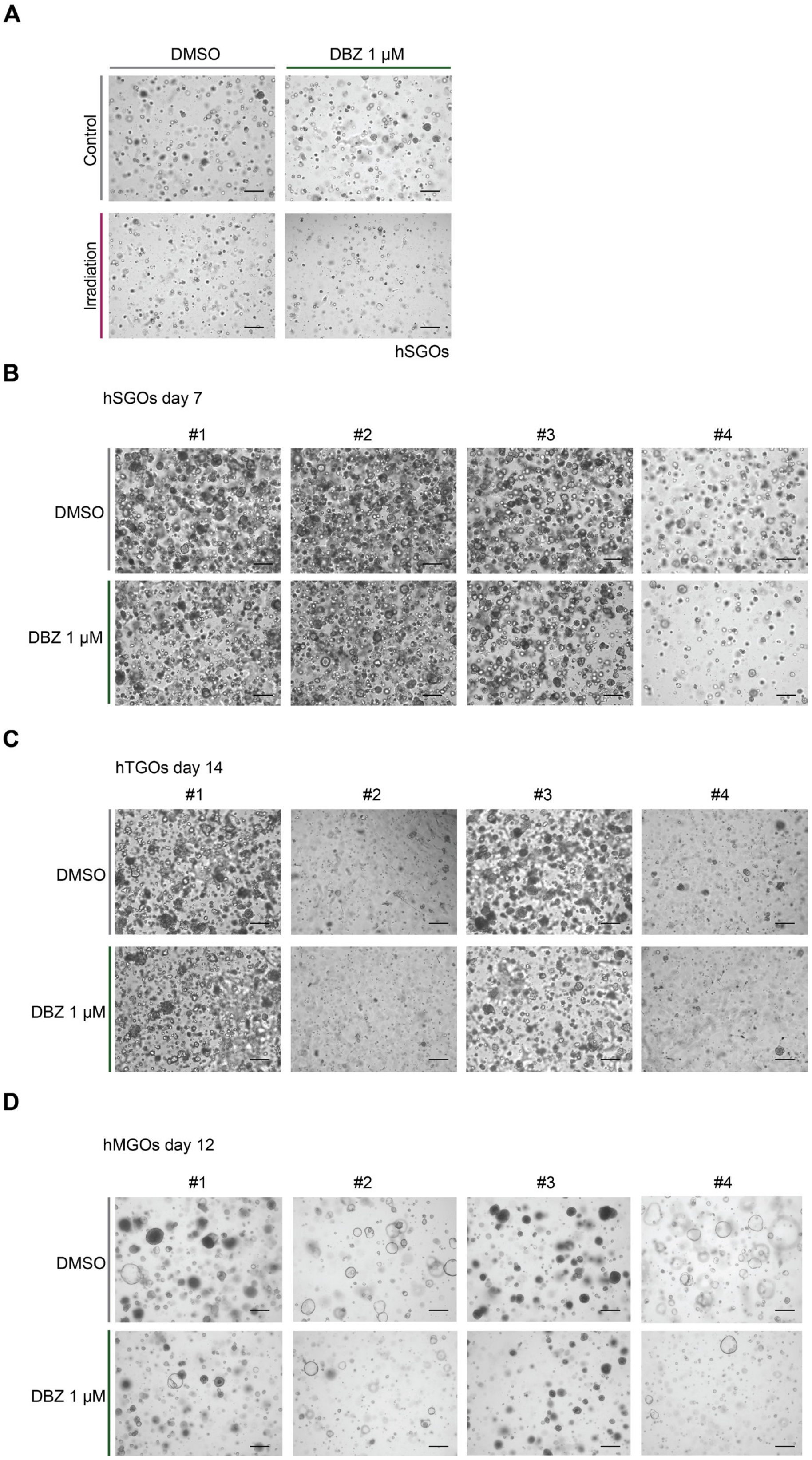


**Figure S6. Patient-derived glandular organoids**

(**A**) Representative images of control and photon-irradiated hSGOs after treatment with DMSO, DBZ or JAG1. (**B**) Representative images of hSGOs after DMSO and DBZ treatment. (**C**) Representative images of hTGOs after DMSO and DBZ treatment. (**D**) Representative images of hMGOs after DMSO and DBZ treatment. In all images numbers indicate different patients. Scale bar, 100 μm.

**Table S1. Details of all reagents and antibodies**

| Reagents and antibodies | Company | Catalog number |
|---|---|---|
| **Salivary gland organoid culture and differentiation** | | |
| DMEM/F12 | Gibco/Invitrogen | 11320-074 |
| Penicillin- streptomycin antibiotics | Invitrogen | 15140-163 |
| Glutamax | ThermoFisher Scientific | 35050038 |
| EGF | Sigma-Aldrich | E9644 |
| FGF2 | Peprotech | 100-18-B |
| N2 | Gibco | 17502-048 |
| Insulin | Sigma-Aldrich | I6634-100MG |
| Dexamethasone | Sigma-Aldrich | d4902-25mg |
| 0.05% trypsin EDTA | Invitrogen | 25300-096 |
| Matrigel | Vwr | 356235 |
| Y27632 | Abcam | ab120129 |
| Noggin | Preprotech | 120-10C |
| A8301 | Tocris Bioscience | 2939 |
| Collagenase type II | Gibco | 17101-015 |
| Hyaluronidase | Sigma | H3506-5G |
| $CaCl_2$ | Sigma | C3306 SIGMA |
| HBSS | Gibco | 14175-129 |
| BSA 7.5% | Gibco | 15260037 |
| Fetal bovine serum | Bodinco | 500 ml |
| HGF | Preprotech | 100-39 |
| **Thyroid gland organoid culture and differentiation** | | |
| B27 supplement | Gibco | 17504001 |
| CTS B27 supplement XenoFree | Gibco | A5047501 |
| R-Spondin-1 human | Sigma | SRP3292-20UG |
| UltraGRO-PURE GI Cell Culture Supplement Xeno-free | Pelobiotech | PB-HPCHXCGLI05 |
| HEPES | Thermo Scientific | 15630080 |
| ITS | Gibco | 41400045 |
| bTSH | Sigma | T8931 |
| FGF10 | PeproTech | 100-26 |
| **Mammary gland organoid culture** | | |
| Advanced DMEM/F12 | Gibco/Invitrogen | 12634028 |
| Nicotinamide | Sigma | N0636 |
| N-acetylcysteine | Sigma-Aldrich | A9165-5G |
| Primocin | InvivoGen | ant-pm-05 |
| Hydrocortisone | Cayman Chemical | 20739 |
| β-estradiol | Sigma-Aldrich | E2257 |
| Forskolin | BioGems | 6652995 |
| Heregulin β1 | PeproTech | 100-03 |
| SB202190 | Cayman Chemicals | 10010399-25 |
| FGF7 | Immunotools | 11343653 |
| Culturex growth factor reduced BME | R&D Systems | 3433-010-01 |
| **Organoid treatments** | | |
| DBZ | Axon Medchem | 1488 |

| | | |
|---|---|---|
| JAG1 | Alpha Diagnostic | SP-56614-1 |
| DMSO | Sigma | 276855-100 ML |
| DAPT | Sigma | D5942-5MG |
| **Florescence activated cell-sorting** | | |
| Propidium iodide | Sigma | P4170 |
| Anti-rat CD29 FITC (1:200) | BD biosciences | 555005 |
| Anti-mouse CD24 PB (1:200) | Biolegend | 101820 |
| Anti-mouse CD44 PE (1:200) | BD biosciences | 553134 |
| **Immunofluorescence staining** | | |
| HistoGel embedding medium | Epredia | HG-4000-012 |
| Tris base | Merck | 10708976001 |
| EDTA | Invitrogen | 15576-028 |
| Donkey serum | Jackson Immuno Research | 017-000-121 |
| Triton X-100 | Sigma | T8787-250ML |
| KRT14 (1:300) | Abcam | ab7800 |
| KRT8/18 (1:50) | DSHB | ab531826 |
| CD29 (1:200) | Cell Signaling | 34971 |
| CD44 (1:200) | Biolegend | 103002 |
| AQP5 (1:300) | Abcam | ab92320 |
| SOX9 (1:200) | Cell Signaling | 82630 |
| CD164 (1:300) | Santa Cruz | sc-271179 |
| NIS (Slc5a5) (1:100) | Proteintech | 24324-1-AP |
| Thyroglobulin (TG) (1:100) | Dako | A0251 |
| DAPI | Sigma Aldrich | D9542 |
| Mounting medium | Dako | S3025/149699 |
| Donkey anti-Rabbit IgG, Alexa Fluor 488 | Invitrogen | A-21206 |
| Donkey anti-Mouse IgG, Alexa Fluor 594 | Invitrogen | A-21203 |
| Donkey anti-Rat IgG, Alexa Fluor 594 | Invitrogen | A-21209 |
| **Quantitative real-time qPCR** | | |
| dNTP mix | Invitrogen | 10297-018 |
| Random primers | Invitrogen | SO142 |
| First-strand Buffer | Invitrogen | 28025013 |
| DTT | Invitrogen | 328025013 |
| RNase OUTTM | Invitrogen | 10777019 |
| M-MLV RT | Invitrogen | 28025013 |
| iQ SYBR Green Supermix | Bio-Rad | 170-8885 |
| **Single cell RNA-sequencing** | | |
| 10X Chromium Next GEM Single Cell 3' Kit v3.1 | 10X Genomics | 1000128 |
| Single Index Kit T Set A | 10X Genomics | 1000213 |
| 10X Chromium Next GEM Chip G Single Cell Kit | 10X Genomics | 1000127 |
| **ATAC-sequencing** | | |
| Nextera DNA Sample Preparation Kit | Illumina | FC-121-1030 |
| Qiagen MinElute kit | Qiagen | 28004 |
| E-gel agarose gel | Thermo Fisher Scientific | G521802 |
| Zymoclean Gel DNA Recovery Kit | Zymo | D4007 |

**Table S2. Primer sequences (mouse)**

| Gene | Forward Primer 5`- 3` | Reverse Primer 5` - 3` |
|---|---|---|
| *Ywhaz* | TTACTTGGCCGAGGTTGCT | TGCTGTGACTGGTCCACAAT |
| *Notch1* | AGCAAGAAGAAGCGGGAGAGC | TGTCGTCCATCAGAGCACCATC |
| *Hes1* | CACTGGAAGGTGACACTGCG | GAGAGGCTGCCAAGGTTTTTG |
| *Jag1* | ACACAGGGATTGCCCACTTC | AGCCAAAGCCATAGTAGTGGTCA |
| *Hey1* | CCCAAACTCCGATAGTCCATAGCC | GCCGACGAGACCGATCAATAAC |
| *Nkx2-1* | CGCCTTACCAGGACACCAT | CCCATGCCACTCATATTCAT |
| *Slc5a5* | TCCACAGGAATCATCTGCACC | CCACGGCCTTCATACCACC |
| *Tpo* | ACAGTCACAGTTCTCCACGGATG | ATCTCTATTGTTGCACGCCCC |
| *Tg* | AGGACCCGTGTGGTAGG | CTGACCCAGAGAATGGCAGT |
